# SK2/3 CHANNELS COUPLE WITH T-TYPE CA^2+^ CHANNELS TO GATE SPINAL LOCOMOTOR RHYTHM GENERATION

**DOI:** 10.64898/2026.03.19.712770

**Authors:** Florent Krust, Cyprien Dautrevaux, Cécile Brocard, Virginie Trouplin, Benoît Drouillas, Jean-Didier Lemaréchal, Meysam Hashemi, Mathieu Gilson, Frédéric Brocard

## Abstract

Initiating locomotion requires central pattern generator (CPG) interneurons to transition from tonic firing to persistent sodium current (I_NaP_)-dependent bursting. While I_NaP_ provides the rhythmogenic drive, the conductances gating this transition remain unclear. Here, we show that functional coupling between small-conductance calcium-activated potassium (SK2/3) channels and low-threshold T-type Ca^2+^ channels, notably Cav3.2, gates locomotor rhythm generation. Pharmacological or genetic disruption of this SK-T-type axis triggers intrinsic bursting in Hb9 interneurons, a genetically identified rhythmogenic population of the locomotor CPG, and initiates fictive locomotion, whereas SK activation silences ongoing rhythmic output. Immunohistochemical co-expression of SK2, SK3 and Cav3.2 in Hb9 interneurons provides an anatomical basis for this functional coupling. Simulation-based inference further shows that, beyond this gating mechanism, burst diversity is primarily determined by the balance between I_NaP_ and M-type potassium conductances. Together, these findings identify SK-T-type coupling as a tunable brake on CPG activation, defining a biophysical module that controls the initiation and termination of locomotor rhythmic activity, with potential relevance for rhythmogenic circuits beyond locomotion.

## Introduction

Locomotor activity arises from spinal circuits organized into a central pattern generator (CPG) capable of producing rhythmic movement in the absence of sensory or supraspinal drive ^1, 2^. At the core of this rhythmogenic network lie glutamatergic interneurons ^3^, including a subset of pacemaker neurons that intrinsically generate membrane oscillations ^4, 5^. These neurons rely on the persistent sodium current (I_NaP_) ^6–9^ whose critical role in locomotor rhythm generation is well established across vertebrate CPGs ^6, 10–15^. However, while the “engine” of these rhythms, I_NaP_, is well-characterized, the mechanisms that gate the transition from silence to rhythmic locomotor output remain incompletely understood.

During locomotor onset, extracellular calcium ([Ca^2+^]_o_) and potassium ([K^+^]_o_) concentrations undergo fluctuations that promote the emergence of pacemaker bursting ^16^. These ionic changes enhance I_NaP_ while reducing outward potassium currents that normally oppose it ^16^. While Nav1.1 and Nav1.6 driving I_NaP_ are well-characterized ^17^, the potassium channels that restrain burst emergence and the subsequent initiation of the locomotor rhythm remain poorly defined. The Kv7.2-mediated M-current (I_M_) contributes to the regulation of locomotor speed ^18^, but it does not fully account for the potassium-dependent brake that limits CPG activation.

Given the dynamic changes in [Ca^2+^]_o_ and [K^+^]_o_ during locomotor onset ^16^, small-conductance calcium-activated (SK) channels are ideally positioned to convert these ionic fluctuations into changes in firing mode. SK channels are classically viewed as burst-terminating conductances that mediate spike-triggered afterhyperpolarization (AHP) and regulate locomotor output ^15, 19–21^. However, emerging evidence suggests a more complex role for these channels in initiating rhythmic activity ^22, 23^, while the relevant isoforms, their subcellular organization, and their calcium triggers within the locomotor CPG remain unknown. Here, through a combination of patch-clamp recordings, isoform-specific viral knockdown, high-resolution immunohistochemistry, and state-of-the-art simulation-based inference, we challenge the classical view of SK channels as simple feedback regulators. We show that functional coupling between SK2/3 channels and T-type Ca^2+^ channels (notably Cav3.2) gates pacemaker bursting and locomotor rhythm initiation. These findings identify an atypical SK-T-type mechanism that may represent a broader biophysical module for gating oscillatory activity across rhythmogenic circuits.

## Results

### Potassium channels constrain I_NaP_-driven burst expression

We first examined whether inhibiting potassium channels unmasks intrinsic bursting in interneurons from the rhythmogenic locomotor CPG region (ventromedial lamina VII-VIII, L_1_-L_2_) by performing whole-cell recordings in spinal slices from neonatal mice (P5-12). In standard conditions (aCSF: [Ca^2+^] = 1.2 mM and [K^+^] = 3 mM), interneurons fired tonically and did not burst ^6, 16^. However, when intracellular K^+^ currents were blocked with a CsCl-based internal solution, bursting emerged in 70% of interneurons (21/30 cells; Fig. 1a). Bursts were characterized as long-lasting plateau-like depolarizations (duration: 5.7 ± 0.9 s; amplitude: 37.9 ± 3.9 mV; area: 171.9 ± 50.2 mV·s), occurring at low frequency (0.2□±□0.03□Hz; Fig. 1b). Following individual spikes, the afterhyperpolarization (AHP) was reduced or absent, often replaced by a slow afterdepolarization (Fig. 1c, arrowhead) which could evolve into plateau potentials upon brief depolarizing steps (Fig. 1d). Tetrodotoxin (TTX; 0.5 µM) abolished bursting and eliminated plateau potentials prior to spike attenuation (Fig. 1d,e), consistent with the involvement of a TTX-sensitive I_NaP_ in this intrinsic bursting activity.

**Figure 1.**
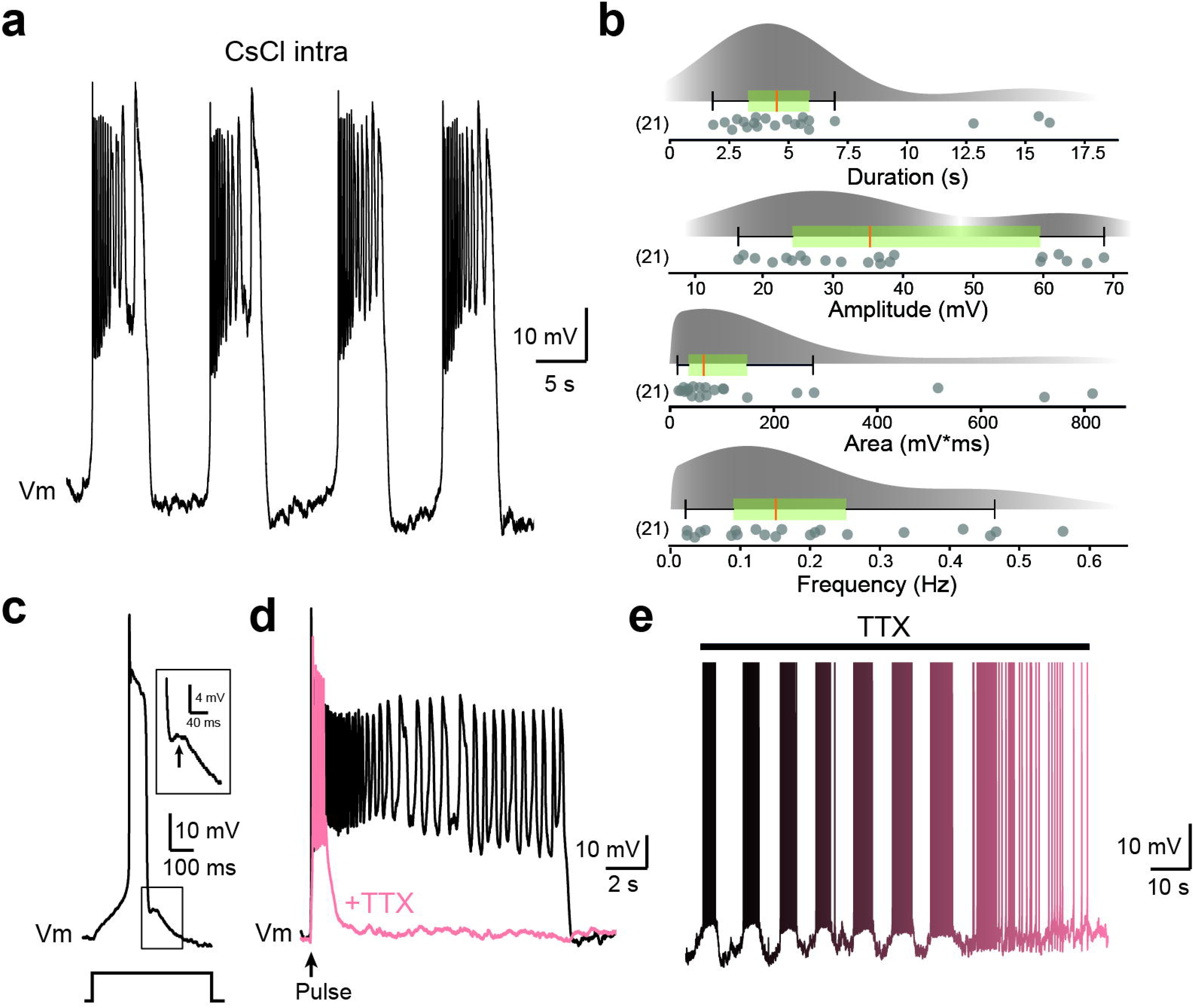
Potassium channels restrain I_NaP_-driven burst expression. **a** Voltage trace from ventromedial interneuron of the rhythmogenic CPG region (L_1_-L_2_) recorded with a CsCl-based intracellular solution, showing rhythmic bursting. **b** Raincloud plots with box-and-whisker overlays (median, interquartile range) quantifying burst duration, amplitude, area, and frequency. Each dot represents a single cell; sample size is indicated in parentheses. **c** Representative action potential evoked by a near-threshold depolarizing pulse. Inset highlights the slow afterdepolarization (ADP, arrow) replacing AHP. **d** Voltage responses to a 2-ms suprathreshold pulse before (black) and after (pink) tetrodotoxin (TTX, 0.5 µM) application. **e** Transition from bursting to tonic firing during TTX wash-in.

### SK channels restrain burst emergence in the rhythmogenic region of the locomotor CPG

Because intracellular Cs^+^ attenuated the AHP and unmasked bursting, we tested whether Ca^2+^-activated K⁺ channels specifically constrain burst initiation. We applied selective blockers of their three main subtypes: BK (IbTx, 200 nM), IK (Tram-34, 5 µM), and SK (apamin, 100-200 nM) channels ^24–26^. While IbTx shortened AHP duration (P < 0.05; Supplementary Fig. 1a,b) without affecting its amplitude (P > 0.05; Supplementary Fig. 1c) and broadened action potentials by slowing repolarization (P < 0.01; Supplementary Fig. 1d-e), effects consistent with BK channel function ^27^, it nonetheless failed to convert tonic spiking into bursting (0/11 cells; Supplementary Fig. 1g,h). Similarly, IK blockade with Tram-34 had no effect on the AHP, spike waveform or firing mode (0/12 cells; Supplementary Fig. 1a-h). These results indicate that while BK channels shape spike repolarization and AHP kinetics, neither BK nor IK channels gate burst initiation.

In contrast, SK channel blockade with apamin induced rhythmic bursting in approximately half of the interneurons (36/73 cells; Fig. 2a-d), an effect accompanied by a reduction in AHP amplitude (P < 0.01; Fig. 2f,g) and duration (P < 0.001; Fig. 2f,h), alongside increased firing frequency (P < 0.001; Fig. 2f,i). This apamin-induced bursting was abolished by riluzole (5 µM), confirming its I_NaP_ dependence (Fig. 2e). Compared to Cs^+^-evoked bursts, those induced by apamin were shorter (P < 0.05; Fig. 2j), displayed smaller amplitude and area (P < 0.001; Fig. 2k,l), yet occurred at a higher frequency (P < 0.001; Fig. 2m). A similar effect was observed in rats, where apamin triggered bursting in 50% of ventromedial interneurons (38/76 cells; Fig. 2b) with burst characteristics comparable to those recorded in mice (P > 0.05; Fig. 2n-q).

**Figure 2.**
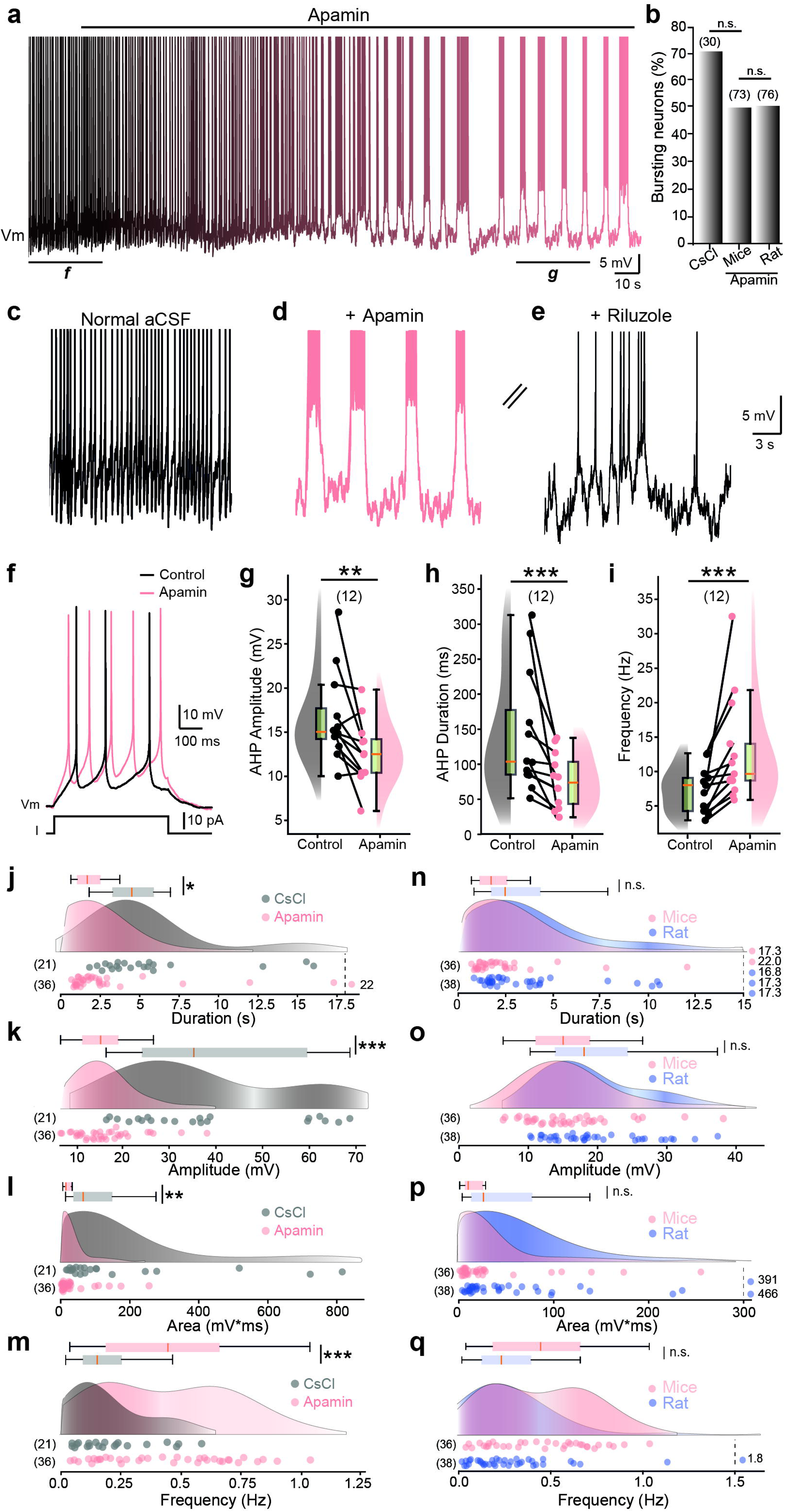
SK channels gate I_NaP_-dependent bursting activity. **a** Example trace showing the transition from tonic firing to rhythmic bursting following apamin application. **b** Bar graph showing the proportion of bursting neurons with CsCl-based intracellular solution or under bath-applied apamin. **c-d** Expanded segments from the same in **a** showing activity in standard aCSF (**c**) and after apamin (**d**). **e** Bursting abolished by riluzole (5 µM). **f** Voltage traces from ventromedial interneurons of the rhythmogenic CPG region (L_1_-L_2_) evoked by near-threshold depolarizing pulses before (black) and after (pink) apamin application (200 nM). **g-i** Raincloud plots with boxplots (median and interquartile range) quantifying afterhyperpolarization (AHP) amplitude (**g**), AHP duration (**h**), and spiking frequency (**i**) before and after apamin. **j-m** Raincloud plots with boxplots (median and interquartile range) quantifying burst duration (**j**), amplitude (**k**), area (**l**), and frequency (**m**) in neurons recorded with CsCl-based intracellular solution (grey) or under bath-applied apamin (pink). **n-q** Raincloud plots with boxplots (median and interquartile range) quantifying burst duration (**n**), amplitude (**o**), area (**p**), and frequency (**q**) after apamin in mouse (pink) and rat (blue) interneurons. Data points plotted beyond the dashed vertical line indicate values outside the axis range. Numbers in parentheses indicate recorded cells; each dot represents a single cell. n.s., not significant; \**P* < 0.05; \*\**P* < 0.01; \*\*\**P* < 0.001 (two-sided Fisher’s exact test for **b**; two-sided Wilcoxon paired test for **g-i**; two-sided unpaired t-test for **j-q**). For detailed *P* values, see Source data.

To further validate SK involvement, we used UCL-1684, a broad-spectrum SK channel blocker ^28, 29^ structurally distinct from apamin ^30^. UCL-1684 induced dose-dependent bursting (Supplementary Fig. 2a-c), reaching ∼50% of bursters at 1 µM (8/15 cells) with features similar to those evoked by apamin (P > 0.05; Supplementary Fig. 2d-g). Together, these findings indicate that SK channels are the main Ca^2+^-activated K^+^ conductance limiting I_NaP_-dependent bursting in interneurons of the spinal locomotor CPG region.

### SK2/3 isoforms gate bursting in CPG interneurons

Although apamin and UCL-1684 unmask rhythmic bursting, their limited selectivity across SK1-3 isoforms precludes precise subunit attribution. In rodents, SK1 is relatively insensitive to these blockers ^31^, suggesting the involvement of SK2 and/or SK3. We therefore applied tamapin, a peptide toxin with high affinity for SK2 and SK3 ^32^. Tamapin induced bursting in a concentration-dependent manner, nearly doubling the fraction of bursting interneurons at 10 nM versus 5 nM and reaching ∼50% of the recorded population (Fig. 3a,b,d). This bursting was accompanied by reduced AHP amplitude (P < 0.01; Fig. 3e,f) and duration (P < 0.05; Fig. 3e,g), alongside increased firing frequency (P < 0.01; Fig. 3e,h). Furthermore, burst parameters matched those evoked by apamin (P > 0.05; Fig. 3i-k). Notably, tamapin also promoted bursting in genetically identified Hb9 interneurons (3/7 cells; Fig. 3d), indicating that SK2/3-dependent bursting also occurs in a genetically identified rhythmogenic population of the locomotor CPG ^6, 33^.

**Figure 3.**
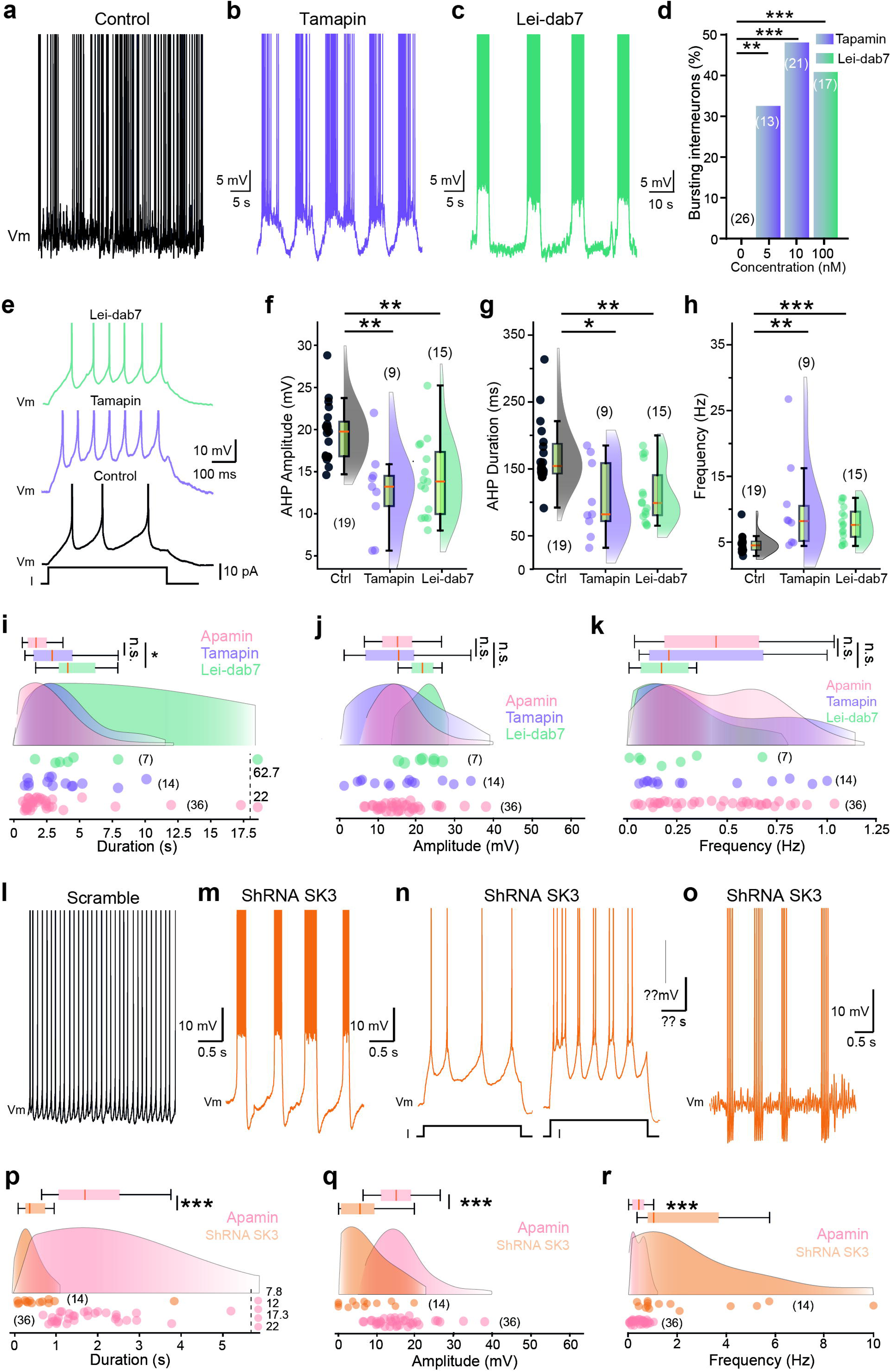
Complementary roles of SK2 and SK3 in burst initiation. **a-c** Voltage traces from ventromedial interneurons of the rhythmogenic CPG region (L1-L2) recorded under control conditions (**a**), after tamapin application (10 nM, **b**), or after lei-dab7 application (10 nM, **c**). **d** Bar graph showing the proportion of interneurons displaying bursting in response to increasing concentrations of lei-dab7 (teal) or tamapin (purple). **e** Representative action potentials evoked by near-threshold current injections under control conditions (black), tamapin (purple), or lei-dab7 (teal). **f-h** Raincloud plots with box-and-whisker overlays (median, interquartile range) quantifying AHP amplitude (**f**), AHP duration (**g**), and firing frequency (**h**) under the three conditions. **i-k** Raincloud plots with box-and-whisker overlays (median, interquartile range) quantifying burst duration (**i**), amplitude (**j**), and frequency (**k**) across interneurons treated with apamin (pink), tamapin (purple), or lei-dab7 (teal). **l-o** Representative voltage traces from interneurons expressing control shRNA (**l**) or SK3-targeting shRNA (**m-o**), showing heterogeneous bursting phenotypes: bursts at rest/rheobase (**m**), bursts requiring stronger depolarization (**n**), and elliptic bursting dynamics (**o**). **p-r** Raincloud plots comparing burst duration (**p**), amplitude (**q**), and frequency (**r**) in interneurons after SK3 knockdown (orange) and after apamin application (pink). Numbers in parentheses denote recorded cells; each dot represents a single cell. Data points plotted beyond the dashed vertical line indicate values outside the axis range. n.s., not significant; \**P* < 0.05; \*\**P* < 0.01; \*\*\**P* < 0.001 (two-sided Fisher’s exact test for **d**; Kruskal-Wallis with Dunn’s post hoc test versus control for **f-h** and **i-k**; two-sided Mann-Whitney test for **p-r**). For detailed *P* values, see Source Data.

To further isolate the role of SK2, we used lei-dab7, a derivative selective for SK2 over SK3 ^34^. Lei-dab7 induced bursting in 41% of interneurons (7/17 cells; Fig. 3c,d) and reproduced the classic SK-blockade signature: AHP amplitude and duration decreased (P < 0.01; Fig. 3e-g) with increased firing frequency (P < 0.001; Fig. 3e,h). Lei-dab7-induced bursts were significantly longer than those evoked by apamin (P < 0.05; Fig. 3i), whereas their amplitude and frequency remained similar (P > 0.05; Fig. 3j,k).

To assess the specific contribution of SK3, for which no selective pharmacological blocker is available, we turned to RNA interference. An SK3-targeting shRNA, validated in HEK-293 cells (∼45% protein reduction; Supplementary Fig. 3a,b), was delivered intrathecally at birth (T_13_-L_1_) via an AAV9 vector. While control (scrambled shRNA) eGFP^+^ interneurons remained tonically active (n = 8 cells; Fig. 3l), SK3 knockdown promoted bursting in ∼42% of cells (14/33 cells). SK3 knockdown triggered bursting without altering AHP properties (P > 0.05; Supplementary Fig. 3c-e). SK3 reduction revealed three bursting profiles: spontaneous bursting at rest (n = 6 cells; Fig. 3m), depolarization-induced bursting (n = 4 cells; Fig. 3n), and an elliptic bursting pattern (n = 4 cells; Fig. 3o). Overall, SK3-knockdown bursts were shorter (P < 0.001; Fig. 3p), smaller (P < 0.01; Fig. 3q), and occurred at a higher frequency (P < 0.001; Fig. 3r) than those triggered by apamin.

Together, these findings identify SK2 and SK3 as the principal SK isoforms constraining I_NaP_-driven bursting in locomotor CPG interneurons.

### Distinct somatodendritic organization of SK2 and SK3 subunits in Hb9 interneurons

To determine whether SK2 and SK3 display distinct subcellular distributions in Hb9 interneurons, we mapped their localization by high-resolution confocal immunofluorescence in eGFP-labeled Hb9 interneurons (Fig. 4a). Both antibodies produced punctate labeling organized into discrete clusters, distributed across somatic and dendritic compartments (Fig. 4b,e,f). At the soma, however, SK2 and SK3 showed distinct distribution patterns. SK2 assembled into prominent membrane-associated clusters outlining the somatic contour (Fig. 4b), whereas SK3 labeling was more diffuse and less enriched at the somatic membrane. In line with this asymmetric distribution, somatic SK2 clusters were significantly larger (P < 0.001; Fig. 4c) and displayed a higher density per unit membrane area (P < 0.001; Fig. 4d). Co-localized SK2/3 clusters were rare at the soma (3.3 %) and occurred at the lowest density compared with SK2- or SK3-only clusters (P < 0.001; Fig. 4d).

**Figure 4.**
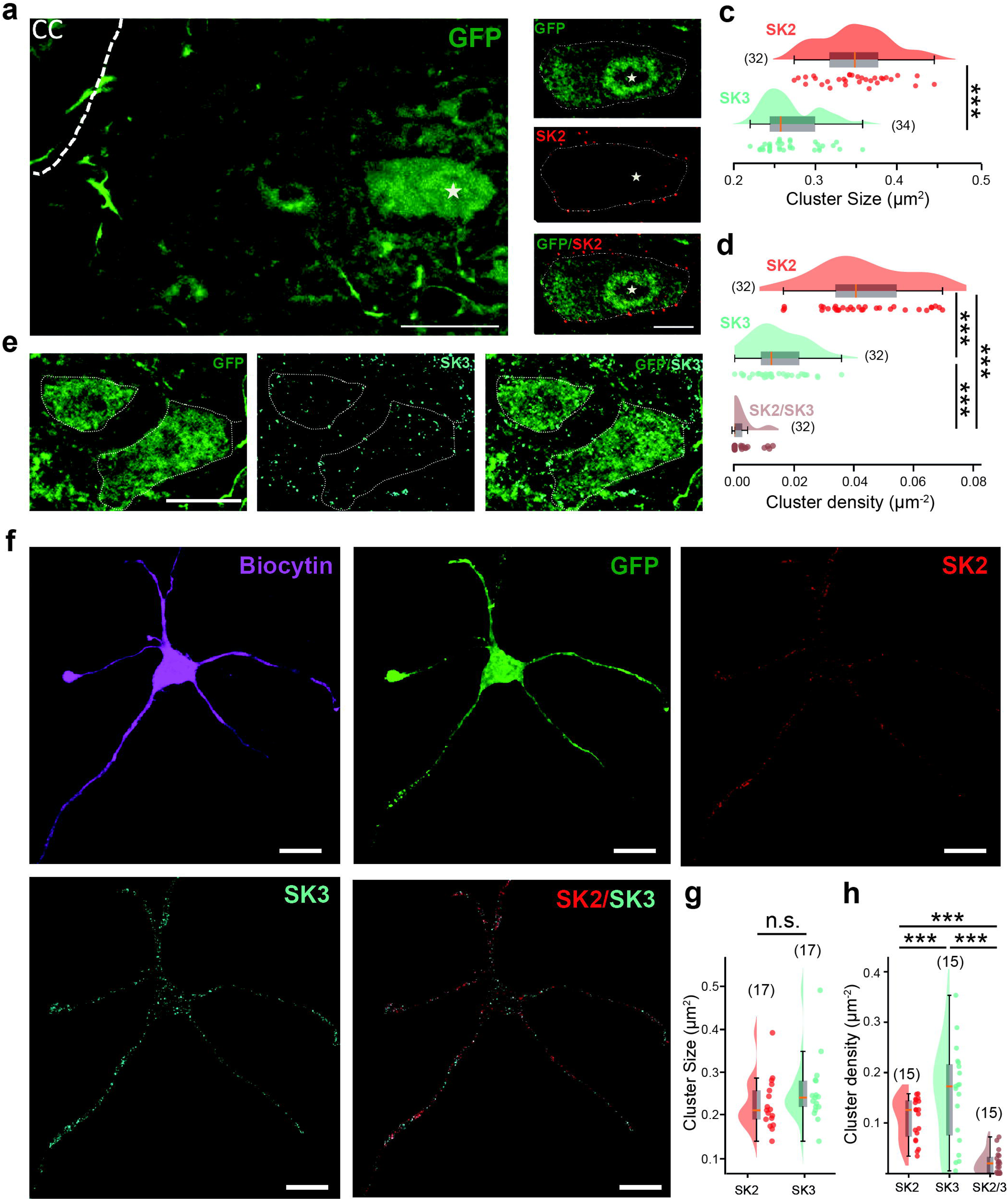
Distribution of SK2 and SK3 channels in Hb9-positive interneurons. **a** Representative low-magnification confocal image of GFP-expressing Hb9 interneurons in the ventromedial spinal cord. The asterisk marks the soma of an identified interneuron. The central canal (cc) is indicated by the dashed line. Scale bar, 25 µm. **b** High-magnification images of the soma indicated in **a**, showing GFP fluorescence (top), SK2 immunolabeling (middle), and a merged GFP/SK2 overlay (bottom). Dashed lines delineate the somatic membrane. Scale bar, 10 µm. **c-d** Raincloud plots with box-and-whisker overlays (median, interquartile range) quantifying the size (**c**) and density (**d**, clusters per µm^2^) of SK2, SK3, and co-localized SK2/3 clusters at the somatic membrane. **e** Confocal images of an Hb9 interneuron soma showing GFP (left), SK3 immunolabeling (middle), and a merged GFP/SK3 overlay (right). Dashed lines delineate the somatic membrane. Scale bar, 20 µm. **f** Confocal images of an intracellularly recorded HB9 interneuron filled with biocytin (magenta), showing GFP immunofluorescence (green), SK2 (red), and SK3 immunolabeling (cyan), alongside a merged GFP/SK3 signal. Scale bar, 20 µm. **g-h** Raincloud plots with box-and-whisker overlays (median, interquartile range) quantifying the size (**g**) and density (**h**) of SK2, SK3, and co-localized clusters along the dendrites. Numbers in parentheses denote the number of cells (**c, d**) or dendrites (**g, h**) analyzed; each dot represents a single measurement. *** *P* < 0.001 (two-sided unpaired t-test for **c** and **g**; One-way ANOVA for **d** and **h**). For detailed *P* values, see Source data.

Because standard immunolabeling does not allow unambiguous attribution of dendritic processes to individual Hb9 interneurons in spinal cord sections, we performed whole-cell recordings with biocytin filling followed by post hoc immunolabeling to expand the analysis to their dendrites (Fig. 4f). This approach revealed intermingled SK2 and SK3 clusters distributed along dendritic shafts (Fig. 4f and Supplementary Fig. 4a,b). In contrast to the soma, SK2 and SK3 cluster sizes were similar in dendrites (P > 0.05; Fig. 4g). However, their densities diverged, with SK3 clusters being significantly more abundant along dendrites than SK2 clusters (P < 0.001; Fig. 4h and Supplementary Fig. 4b). Co-localized SK2/3 clusters again displayed the lowest density (P < 0.05; Fig. 4h), representing 17% of total dendritic clusters.

Together, these data reveal a clear spatial segregation of SK subunits in Hb9 interneurons, with SK2 enriched at the soma and SK3 predominating along dendrites.

### SK channels bidirectionally gate rhythmic bursting dynamics

We next asked whether SK channels also shape the dynamics of ongoing oscillations once bursting is established. Under locomotor-like ionic conditions ([Ca^2+^]_o_ = 0.9 mM; [K^+^]_o_ = 6 mM), which elicit bursting in ∼50% of CPG interneurons ^16^, apamin potentiated burst output (Fig. 5a). Specifically, it increased intraburst spike frequency (P < 0.01; 21.2 ± 3.7 Hz vs 49.2 ± 8.9 Hz, n = 10 cells), as well as burst amplitude and duration (P < 0.01; Fig. 5b,c) while decreasing burst frequency (P < 0.05; Fig. 5d). These results indicate that SK channels limit burst strength and duration while contributing to the pacing of rhythmic activity once bursting is established.

**Figure 5.**
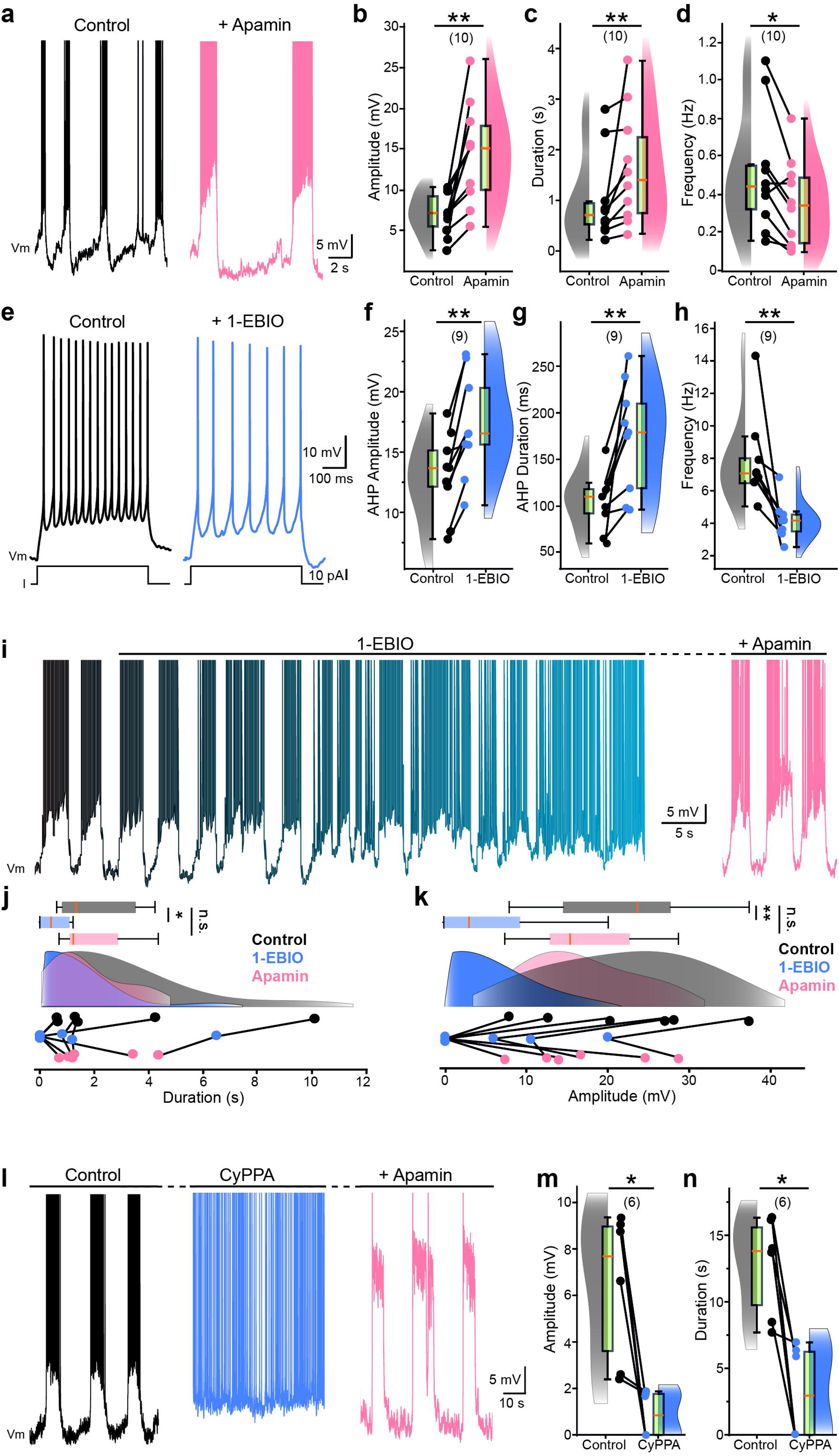
SK channels bidirectionally regulate intrinsic bursting dynamics. **a** Voltage traces from a bursting ventromedial interneuron (L_1_-L_2_) recorded under locomotor-like ionic conditions ([Ca^2+^]□ = 0.9 mM; [K^+^]□ = 6 mM) before (black) and after apamin application (pink). **b-d** Raincloud plots with box-and-whisker overlays (median, interquartile range) quantifying burst amplitude (**b**), duration (**c**), and frequency (**d**) before and after apamin. **e** Voltage responses to a suprathreshold depolarizing current step recorded in standard aCSF ([Ca^2+^]□ = 1.2 mM and [K^+^]□ = 3 mM) under control conditions (black) and after 1-EBIO application (blue). **f-h** Raincloud plots quantifying AHP amplitude (**f**), AHP duration (**g**), and firing frequency (**h**) before and after 1-EBIO. **i** Transition from rhythmic bursting to tonic firing during 1-EBIO application (blue gradient), followed by the restoration of bursting upon apamin co-application (pink) in the same cell. **j, k** Raincloud plots comparing burst duration (**j**) and amplitude (**k**) across the three conditions (Control, 1-EBIO, Apamin). **l** Voltage traces under locomotor-like conditions showing baseline bursting (black), transition to tonic firing after CyPPA application (blue), and recovery of bursting with apamin (pink). **m, n** Raincloud plots quantifying burst amplitude (**m**) and duration (**n**) under control and CyPPA conditions. Numbers in parentheses denote recorded cells; each dot represents a single cell. n.s., not significant; \**P* < 0.05; \*\**P* < 0.01 (two-sided Wilcoxon paired test for **b-d**, **f-h**, **m** and **n**; Friedman test with Dunn’s post hoc for **j** and **k**). For detailed *P* values, see Source data.

To determine whether increasing SK activity produced opposite effects, we applied 1-EBIO (200 µM), a positive allosteric modulator that left-shifts the Ca^2+^-activation curve of SK and IK channels ^35^. In standard aCSF, 1-EBIO increased AHP amplitude and duration while reducing firing frequency (P < 0.01; Fig. 5e-h). Under locomotor-like conditions, 1-EBIO strongly reduced bursting: some cells transitioned to tonic firing (Fig. 5i), whereas others exhibited reduced burst duration (P < 0.05; Fig. 5j) and amplitude (P < 0.01; Fig. 5k). Co-application of apamin (200 nM) restored bursting to control levels (P > 0.05 vs. control; Fig. 5i-k), demonstrating that the effects of 1-EBIO are largely mediated by SK rather than IK channels. SK modulation produced comparable bidirectional effects on I_NaP_-dependent bursting evoked by veratridine in standard aCSF (Supplementary Fig. 5). Specifically, apamin enhanced burst amplitude and duration (P < 0.05; Supplementary Fig. 5a,b), whereas 1-EBIO reduced both (P < 0.05; Supplementary Fig. 5c,d). Together, these results indicate a bidirectional control of I_NaP_-driven rhythmicity by SK channels at the single-cell level.

To refine the relevant SK isoforms, we applied CyPPA (3 µM), a selective positive modulator of SK2/3 with negligible effects on SK1 or IK ^35, 36^. CyPPA similarly suppressed bursting, promoting a burst-to-tonic conversion (P < 0.05; Fig. 5l-n). Subsequent application of apamin reinstated bursting (n = 2; Fig. 5l), supporting the conclusion that SK2/3 channels bidirectionally regulate burst propensity in locomotor CPG interneurons, capable of both initiating and terminating rhythmic output depending on their activity level.

### SK2/3-mediated burst gating depends on T-type Ca^2+^ channel coupling

To identify the Ca^2+^ source activating SK2/3 channels in CPG interneurons, we first blocked voltage-gated Ca^2+^ entry with cadmium (100 µM). Cadmium induced rhythmic bursting in 52 % of interneurons (n = 11/21 cells), with burst duration, amplitude, and frequency similar to those observed with apamin (P > 0.05; Fig. 6a-e). Thus, blocking Ca^2+^ influx mimics SK blockade, suggesting that tonic Ca^2+^ influx normally sustains SK activation to restrain bursting.

**Figure 6.**
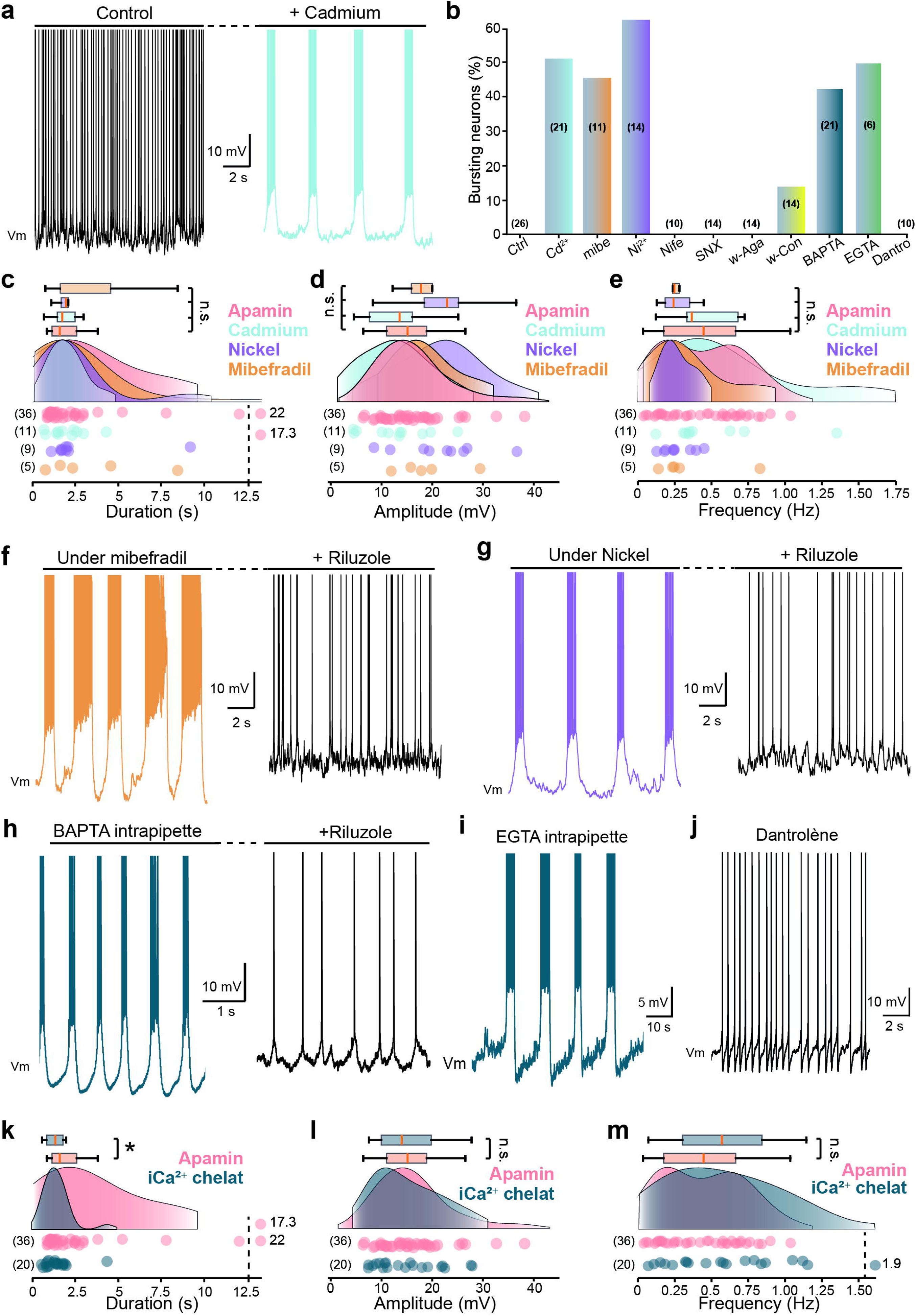
T-type Ca^2+^ channels couple to SK channels to gate intrinsic bursting. **a** Voltage traces from a ventromedial interneuron (L_1_-L_2_) recorded in normal aCSF (black) and after cadmium (Cd^2+^, 100 µM, light blue) application. **b** Bar graph showing the proportion of bursting interneurons under the indicated pharmacological conditions. **c-e** Raincloud plots with box-and-whisker overlays (median, interquartile range) quantifying burst duration (**c**), amplitude (**d**), and frequency (**e**) induced by apamin, Cd^2+^, Ni^2+^, and mibefradil. **f, g** Voltage traces showing rhythmic bursting induced by mibefradil (10 µM, orange, **f**) or nickel (Ni²⁺, 200 µM, purple, **g**) followed by the transition to tonic firing after riluzole application (5 µM, black). **h, i** Voltage traces from neurons recorded with intracellular BAPTA (10 mM; teal, **h**) or EGTA (10 mM; teal, **i**). Note the transition to tonic firing after riluzole application in **h**. **j** Representative voltage trace showing the maintenance of tonic firing (absence of bursting) in the presence of dantrolene (50 µM). **k-m** Raincloud plots with box-and-whisker overlays comparing burst duration (**k**), amplitude (**l**), and frequency (**m**) between chelation-induced bursts (BAPTA or EGTA) and apamin-induced bursts. Numbers in parentheses denote recorded cells; each dot represents a single cell. Data points plotted beyond the dashed vertical line indicate values outside the axis range. n.s., not significant; \**P* < 0.05; \*\**P* < 0.01; \*\*\**P* < 0.001 (Kruskal-Wallis with Dunn’s post hoc for **c-e**; two-sided Mann-Whitney test for **k-m**). For detailed *P* values, see Source data.

We next examined which Ca^2+^ channel subtypes contribute to this effect. Pharmacological blockade of high-voltage-activated (HVA) channels, including L-type (nifedipine, 20 µM; n = 10 cells), R-type (SNX-482, 500 nM; n = 14 cells), or P/Q-type (ω-agatoxin-IVA, 400 nM; n = 14 cells), failed to induce bursting (Fig. 6b). Among HVA blockers, only ω-conotoxin MVIIC (1 µM; P/Q + N-type) triggered bursting in a small subset of cells (14%, n = 2/14), a significantly lower proportion than observed with apamin or cadmium (P < 0.05; Fig. 6b), suggesting at most a minor contribution from N-type channels. In contrast, inhibiting low-voltage-activated (LVA) T-type channels robustly unmasked bursting. Mibefradil (10 µM) or nickel (Ni^2+^; 200 µM) elicited bursting in 45% (n = 5/11 cells) and 64% (n = 9/14 cells) of interneurons, respectively (Fig. 6b,f,g). These bursts were indistinguishable from those induced by apamin or cadmium (P > 0.05; Fig. 6c-e). A submaximal Ni^2+^ concentration (100 µM) was sufficient to reduce AHP amplitude and duration (P < 0.01; Supplementary Fig. 6a,b), and induced bursting in a subset of interneurons (10 %; 2/21 cells; Supplementary Fig. 6c). Consistent with this pharmacological profile, Cav3.2, the most Ni^2+^-sensitive T-type isoform^37^, was detected as discrete somatic clusters on Hb9 interneurons (Supplementary Fig. 6d,e), providing an anatomical substrate for its functional coupling with SK channels. Notably, riluzole abolished bursting evoked by T-type blockers, confirming its I_NaP_ dependence (Fig. 6f,g).

We then investigated whether SK activation depends on spatially restricted Ca^2+^ signaling. Both the fast Ca^2+^ chelator BAPTA (10 mM) and the slower chelator EGTA (10 mM) induced riluzole-sensitive bursting in 44% (7/21 cells) and 50% (3/6 cells) of interneurons, respectively (Fig. 6b,h,i). Chelation-induced bursts were shorter than apamin-evoked bursts (P < 0.05; Fig. 6k), but had similar amplitude and frequency (P > 0.05; Fig. 6l,m), consistent with SK activation within spatially restricted Ca^2+^ microdomains in close proximity to T-type channels. Finally, blocking ryanodine (dantrolene; 50 µM) or IP_3_ receptors (xestospongin C; 2.5 µM) failed to induce bursting (Fig. 6j), arguing against a major contribution from intracellular Ca^2+^ stores.

Together, these results identify T-type channels, notably Cav3.2, as the principal Ca^2+^ source mediating SK2/3 activation and thereby gating intrinsic bursting in locomotor CPG interneurons.

### Apamin unmasks a spectrum of intrinsic oscillatory phenotypes

Apamin revealed heterogeneous intrinsic responses rather than a uniform bursting response. Regardless of the concentration used, approximately half of the interneurons switched to rhythmic bursting, while the others remained tonic with increased firing rates (Fig. 7a). Within the bursting population, burst waveforms were highly heterogeneous, prompting a systematic characterization of this variability. We applied principal component analysis (PCA) followed by hierarchical clustering to map this heterogeneity. The first two components explained 81% of the total variance (PC1: 48%; PC2: 33%), with PC1 primarily capturing waveform properties (amplitude and slopes) and PC2 tracking temporal dynamics (duration and frequency; Fig. 7b,c).

**Figure 7.**
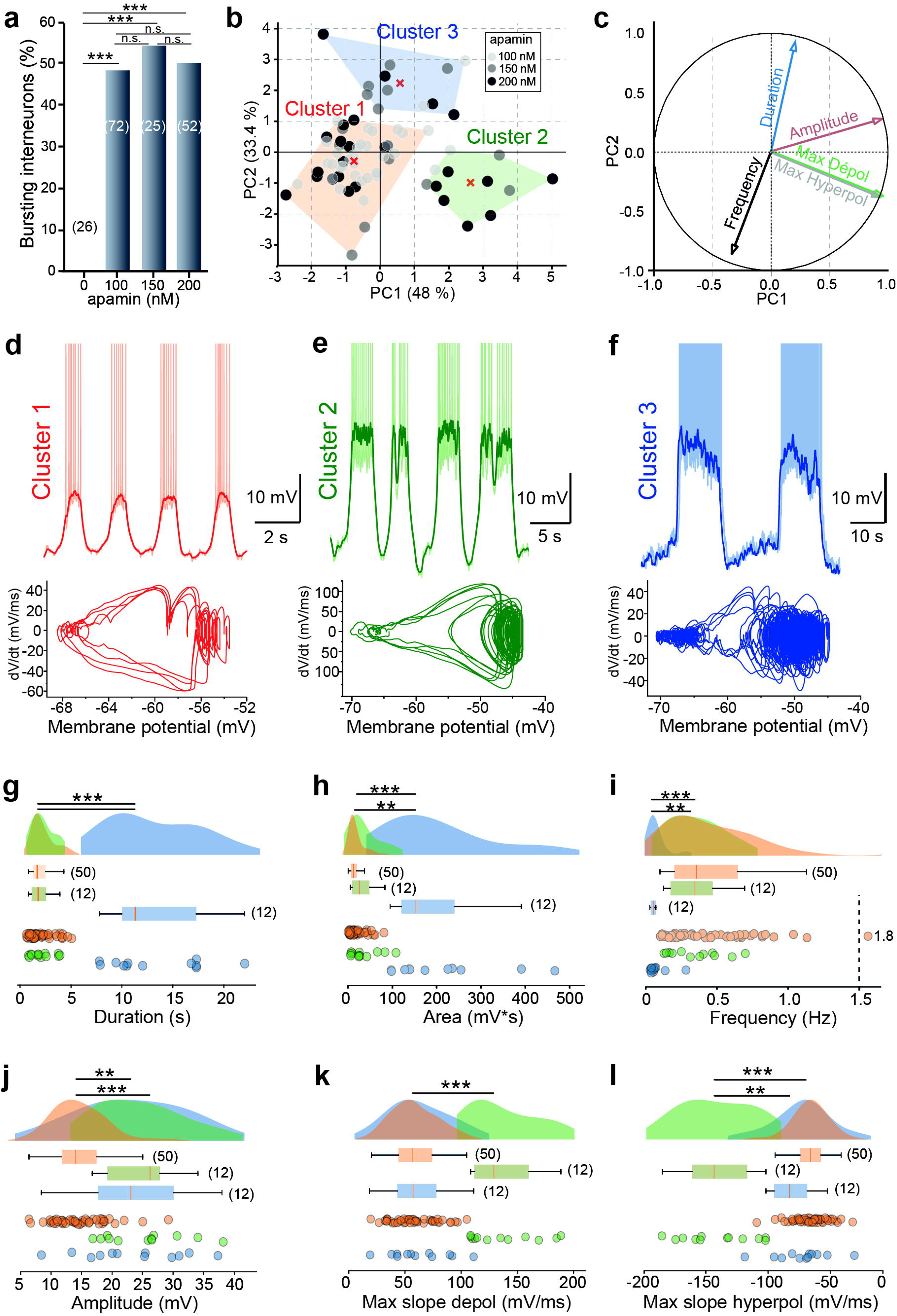
Diversity and biophysical classification of apamin-induced burst patterns. **a** Bar graph showing the proportion of interneurons displaying bursting in response to increasing concentrations of apamin. **b** Principal component analysis (PCA) of burst parameters (duration, amplitude, area, frequency, depolarizing and repolarizing slopes). Hierarchical clustering identified three discrete phenotypes: Cluster 1 (orange), Cluster 2 (green), and Cluster 3 (blue). Red crosses indicate cluster centroids. **c** Correlation circle showing the contribution of each burst parameter to PC1 and PC2. **d-f** Representative recordings for Cluster 1 (**d**), Cluster 2 (**e**), and Cluster 3 (**f**). *Top*: voltage traces showing raw signals with action potentials (shaded) and low-pass filtered signals highlighting burst envelopes (solid lines). Note the different time scales. *Bottom*: corresponding phase-plane trajectories (dV/dt vs voltage). **g-l** Raincloud plots with box-and-whisker overlays (median, interquartile range) showing burst duration (**g**), area (**h**), frequency (**i**), amplitude (**j**), maximal depolarizing slope (**k**), and maximal repolarizing slope (**l**) across the three clusters. Numbers in parentheses denote recorded cells; each dot represents a single cell. Data points plotted beyond the dashed vertical line indicate values outside the axis range. \**P* < 0.05; \*\*\**P* < 0.001 (two-sided Fisher’s exact test for **a**; Kruskal-Wallis with Dunn’s post hoc test for **g-l**). For detailed *P* values, see Source data.

Hierarchical clustering in PCA space (Supplementary Fig. 7a) identified three distinct oscillatory phenotypes (Silhouette score = 0.43; Clusters 1-3; Fig. 7d,f). Cluster 1 (orange) and Cluster 2 (green) shared similar temporal dynamics (duration, area, frequency; P > 0.05; Fig. 7g-i), but differed in waveform. Cluster 1 exhibited the smallest amplitudes (P < 0.01 to P < 0.001; Fig. 7d,j), whereas Cluster 2 displayed the steepest depolarizing and repolarizing slopes (P < 0.01; Fig. 7e,k,l). Cluster 3 (blue) represented a distinct phenotype characterized by slow, plateau-like bursts with the longest durations and lowest frequencies (P < 0.01; Fig. 7f,g-i). While Cluster 3 matched Cluster 2 in amplitude, its slope kinetics more closely resembled the slower dynamics of Cluster 1 (Fig. 7j-l). Together, these comparisons define three distinct oscillatory modes: small/rapid (Cluster 1), large/sharp (Cluster 2), and slow/plateau-like bursts (Cluster 3), each with a characteristic waveform and temporal signature.

Notably, the apamin concentration did not alter the cell distribution in PCA space (P > 0.05; Supplementary Fig. 7b), nor did it affect either principal component axis (P > 0.05; Supplementary Fig. 7c,d). This indicates that the identified phenotypes reflect intrinsic cellular variability rather than dose-dependent effects of SK blockade, prompting us to investigate the conductance combinations associated with each oscillatory mode.

### Simulation-based inference reveals distinct ionic signatures of bursting phenotypes

To uncover the biophysical basis of burst diversity, we used simulation-based inference (SBI) to fit an extended Hodgkin-Huxley model ^16, 18^ to the three experimental phenotypes. The model included SK (g_SK_) and T-type calcium (g_CaT_) conductances (Table 1), with g_SK_ fixed at zero to mimic apamin application. Inference was constrained by summary features extracted from the burst envelope (Supplementary Fig. 8a).

While an initial six-conductance model captured Clusters 1 and 2 phenotypes, it failed to reproduce the slow/plateau dynamics of Cluster 3 (Supplementary Fig. 8b). Because slowing I_NaP_ deinactivation prolongs bursts ^6^, we allowed the maximal I_NaP_ deinactivation time constant (τ_hNaPmax_) to vary as a free parameter. This resolved the mismatch and yielded accurate fits for all three phenotypes (Fig. 8a,b and Supplementary Fig. 8a). Posterior distributions (Fig. 8c) indicated that persistent sodium (g_NaP_) and M-type potassium (g_M_) conductances were the primary determinants of cluster separation. Pairwise posterior distributions further revealed a consistent positive correlation between these conductances across all clusters (Supplementary Fig. 9). The g_NaP_/g_M_ ratio was significantly lower in Cluster 1 than in Clusters 2 and 3 (0.62 vs. 0.93; P < 0.001). Although Clusters 2 and 3 shared a similar ratio (P > 0.05; Fig. 8c), they differed in their absolute conductance regimes (P < 0.001; Fig. 8c), with Cluster 3 additionally requiring increased τ_hNaPmax_ to reproduce its slow/plateau dynamics (Supplementary Fig. 8a). By contrast, g_CaT_ showed broad posterior with no significant differences across the three clusters (P > 0.05; Fig. 8c), indicating a limited contribution to phenotype separation. The remaining conductances, including g_K_, g_Na_ and g_L_, also varied across clusters, although to a lesser extent (Fig. 8c).

**Figure 8.**
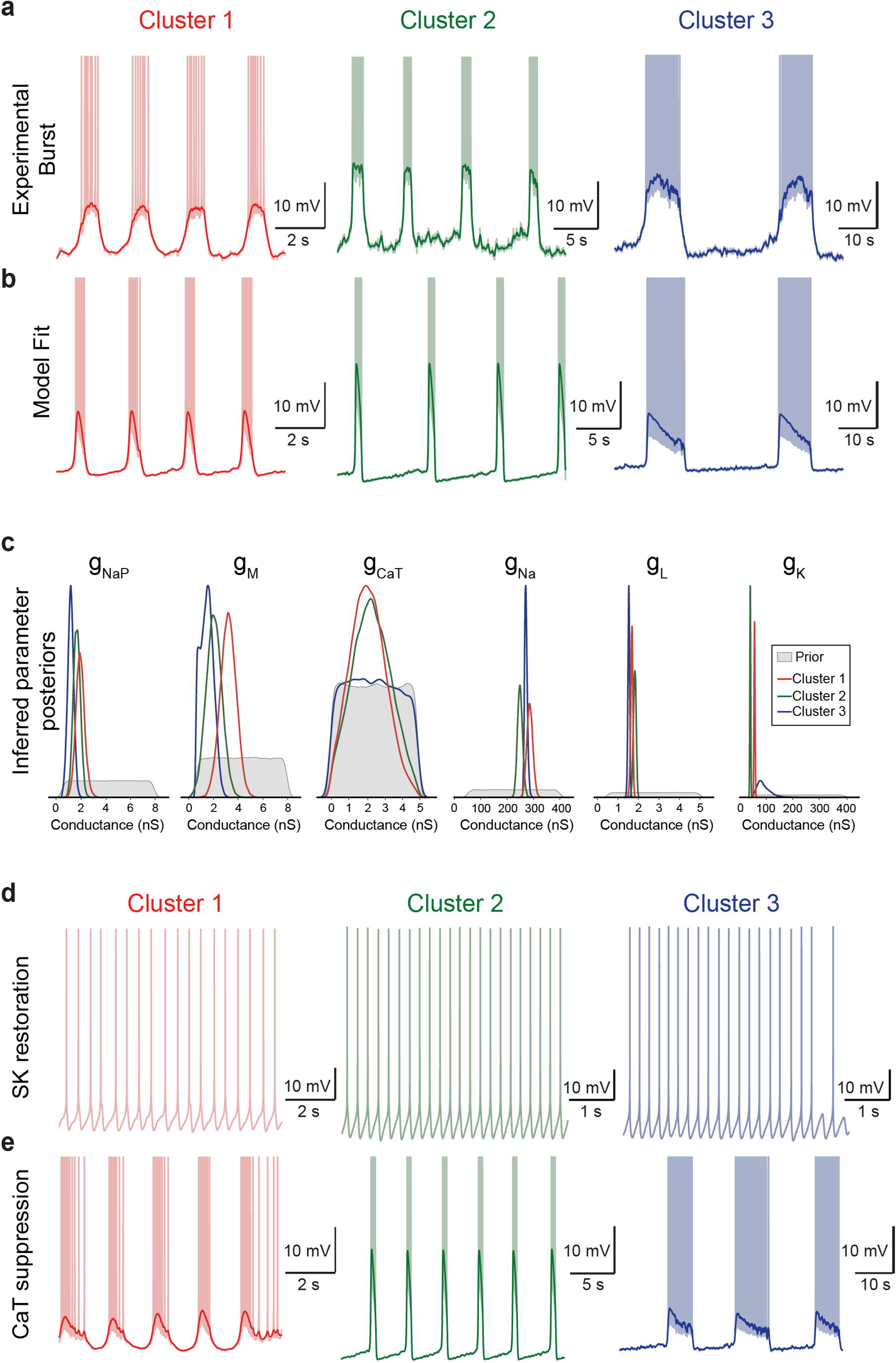
Simulation-based inference identifies the ionic basis of bursting phenotypes. **a** Representative experimental burst for Cluster 1 (red), Cluster 2 (green), and Cluster 3 (blue). Light curves: raw voltage traces; dark lines: burst envelopes used for summary feature extraction. **b** Representative model fits for each phenotype generated from the inferred parameter sets. **c** Posterior distributions of six inferred conductances for each cluster (colored curves) overlaid on the prior distribution (grey shading). **d-e** *In silico* pharmacological manipulations applied to fitted models, consisting of reintroducing SK conductance in **d** (values in Table 2) followed by T-type calcium channel suppression (g_CaT_ = 0) in **e**. For detailed *P* values, see Source data.

*In silico* pharmacological perturbations confirmed the predictive validity of the model. Using cluster-specific parameter values (Table 2), reintroducing g_SK_ converted bursting to tonic spiking across all clusters (Fig. 8d), while subsequent g_CaT_ suppression restored bursting (Fig. 8e), supporting T-type calcium channels as the principal Ca^2+^ source driving SK activation. Suppressing g_NaP_ abolished bursting regardless of cluster identity (Supplementary Fig. 10a), and mimicking EGTA chelation under the SK-on condition reinstated it (Supplementary Fig. 10b), in agreement with the experimental results.

Together, these simulations confirm the predictive validity of the inferred parameters and establish a functional hierarchy in which T-type-SK coupling gates burst expression, I_NaP_ provides the core depolarizing drive, and phenotypic diversity emerges from the balance between g_NaP_ and g_M_, further shaped by I_NaP_ kinetics.

### SK2/3 and T-type channel coupling bidirectionally modulates the onset and termination of locomotor rhythms

We next examined whether the SK2/3-T-type coupling identified at the cellular level scales up to the network level to control locomotor rhythm generation. We recorded fictive locomotion from bilateral L_5_ ventral roots in isolated neonatal mouse spinal cords while applying drugs focally to the rhythmogenic L_1_-L_2_ segments (Fig. 9a-d).

**Figure 9.**
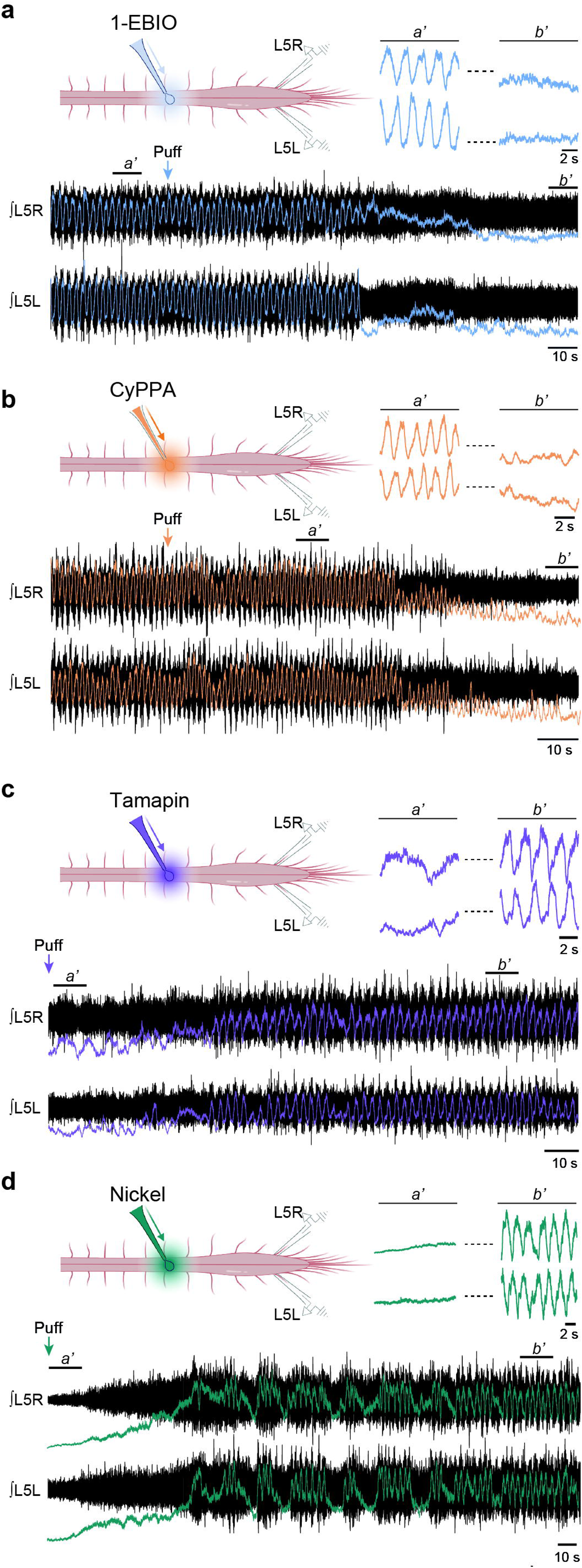
SK2/3 and T-type channel modulation gates locomotor rhythmogenesis. **a-b** Representative ventral root recordings showing the effect of the broad-spectrum SK activator 1-EBIO (**a**; 5 mM) and the SK2/3-selective activator CyPPA (**b**; 0.3-0.4 mM) on ongoing fictive locomotion induced by NMDA/5-HT (5/10 µM). *Top left in* **a** and **b**: Schematic of the isolated spinal cord preparation illustrating focal drug puff application onto rhythmogenic segments (L_1_-L_2_) and bilateral L5 ventral root recordings (L5R, L5L). *Bottom:* Raw ventral root activity (black) superimposed with low-pass filtered envelopes (blue for 1-EBIO, orange for CyPPA); vertical arrows indicate puff onset. *Top right:* Expanded time windows (**a’**, **b’**) of the recordings shown below, highlighting rhythm suppression. **c-d** Representative recordings showing the induction of locomotor-like activity in quiescent preparations (subliminal NMDA/5-HT: 0-1/10 µM) by the SK2/3 inhibitor tamapin (**c**; 1 µM) or the T-type calcium channel blocker nickel (**d**; 10-20 µM). The experimental layout and display are identical to **a, b**. Low-pass filtered envelopes are shown in purple (tamapin) and teal (nickel). Arrows indicate puff onset.

During ongoing fictive locomotion induced by NMDA/5-HT (5/10 µM), focal SK activation with 1-EBIO (5 mM) rapidly and reversibly silenced locomotor output in all preparations (n = 9; Fig. 9a). Similarly, the SK2/3-selective activator CyPPA (0.3-0.4 mM), produced comparable suppression (n = 7; Fig. 9b), indicating that SK2/3 activation is sufficient to terminate ongoing locomotor-like rhythms.

Conversely, we tested whether SK2/3 inhibition could trigger rhythmicity. Under subthreshold NMDA/5-HT conditions (0-1/10 µM), insufficient on their own to generate locomotor output, focal application of tamapin (1 µM) reliably induced locomotor-like activity in all preparations tested (n = 7; Fig. 9c). Consistent with our cellular findings, focal application of the T-type channel blocker Ni^2+^ (10-20 µM) mimicked SK2/3 inhibition and likewise elicited locomotor-like activity (n = 7; Fig. 9d).

Together, these results indicate that the SK2/3-T-type axis operated from single neurons to the CPG network. Inhibition of either component initiates locomotor-like activity, whereas SK2/3 activation terminates ongoing rhythmic output.

## Discussion

Our study establishes that SK channels and T-type Ca^2+^ channels are functionally coupled to gate locomotor rhythm generation in the hindlimb CPG. Converging pharmacological, genetic, immunohistochemical and computational evidence shows that inhibiting SK channels, or blocking T-type Ca^2+^ influx that recruits them, converts tonically firing interneurons into bursters and facilitates rhythmic locomotor output. Conversely, enhancing SK activity suppresses bursting and terminates ongoing locomotor rhythms.

The unmasking of bursting by SK blockade has been reported in a broad range of mammalian brain regions ^38–49^, as well as in invertebrate CPGs ^50^. In these systems, SK currents constrain excitability, such that their removal allows persistent inward currents to dominate and promote rhythmic bursting ^38, 46, 49, 51, 52^. Our recordings support this general framework, showing that apamin-induced bursting depends on I_NaP_, in agreement with our previous work ^4–6, 16–18^. In this respect, the mammalian spinal locomotor CPG appears to follow the same general principle of SK-dependent rhythm control.

We identify SK2 and SK3 as the principal isoforms involved. Their expression in two major locomotor CPG populations^53, 54^, namely Hb9 interneurons as shown here, and Shox2 interneurons as reported previously ^55^, positions them as key regulators of rhythm generation. Whereas all Hb9 interneurons co-express SK2 and SK3, only a subset of Shox2 interneurons expresses both isoforms ^55^, suggesting that distinct rhythmogenic populations may rely on partially different SK-dependent regulatory strategies. This view is further supported by evidence that SK channels contribute to multiple oscillatory regimes. In NMDA-driven bursting, they shape both burst initiation and termination ^56–60^, paralleling what we observe here for I_NaP_-dependent bursting. Given that Hb9 interneurons can alternate between these two modes ^4^, SK channels are well positioned to constrain both regimes in rhythmogenic CPG neurons.

Our data indicate that SK2 and SK3 are not simply redundant. Immunohistochemistry revealed a clear somatodendritic segregation, with SK2 enriched at the soma and SK3 predominating in dendrites. Although SK2/SK3 heteromers have been described in heterologous systems ^61, 62^, the limited SK2/SK3 co-localization observed in our double immunolabeling, together with the lack of co-immunoprecipitation in rat brain ^63^, argues for predominantly homomeric assemblies in situ. Functionally, reducing either isoform promoted bursting, but SK2 inhibition reduced mAHP amplitude whereas SK3 knockdown did not. This difference is consistent with the stronger somatic enrichment of SK2, which would be expected to contribute more directly to the mAHP recorded at the soma, in line with previous evidence that SK2 loss is sufficient to alter the mAHP ^64^ and with the broader view that somatic SK channels are major determinants of mAHP amplitude ^65^. By contrast, the predominantly dendritic localization of SK3 may make its contribution less apparent in this somatic readout. This functional dissociation is also consistent with earlier evidence for isoform-specific roles of SK channels in burst regulation in midbrain dopamine neurons ^66–69^, and with observations in spinal motoneurons, where somatic SK2 regulates baseline AHPs while SK3 predominance in slow-type motoneurons prolongs the AHP and limits firing rates^70, 71^.

Our data further identify transmembrane Ca^2+^ influx, rather than intracellular Ca^2+^ stores, as the main source of SK activation in locomotor CPG interneurons. The promotion of bursting by calcium chelators, together with the lack of effect of intracellular store inhibition, argues that SK recruitment depends primarily on voltage-dependent Ca^2+^ entry. Pharmacological dissection points to T-type Ca^2+^ channels as the dominant source of this influx, since nickel and mibefradil both reduced AHPs and triggered bursting, whereas blockade of high-voltage-activated Ca^2+^ channels was largely ineffective. This interpretation is supported by previous evidence for T-type currents in locomotor-related interneurons ^55^ and by reports that T-type antagonists slow locomotor rhythms ^16, 72^. Interestingly, the effects of T-type channel blockade closely parallel those of SK inhibition, both in locomotor network activity ^19–21, 73^ and in the bursting dynamics of CPG interneurons observed here. Together, these observations strongly support a functional coupling between T-type Ca^2+^ channels and SK channels. Among T-type channel subtypes, Cav3.2 appears as a plausible partner, as we confirm its expression in Hb9 interneurons, consistent with previous work ^72^, and because the high nickel sensitivity we observe is compatible with its known pharmacological profile ^37, 74^.

Computational modeling further refined the hierarchy of mechanisms underlying burst generation and diversification. Although T-type Ca^2+^ channels are essential for recruiting SK current and thereby gating burst expression, simulation-based inference indicates that they do not primarily account for burst diversity. Instead, the balance between g_NaP_ and g_M_ emerged as the main axis separating burst clusters, consistent with earlier work showing that the I_NaP_/I_M_ ratio is a critical determinant of bursting propensity ^18^. Reproducing the slow/plateau phenotype further required adjustment of I_NaP_ deinactivation kinetics, in line with evidence that veratridine prolongs burst duration by slowing I_NaP_ deinactivation ^6^. In this framework, T-type-SK coupling determines whether bursting is permitted, whereas the burst phenotype is shaped primarily by the balance between I_NaP_ and I_M_, further modulated by I_NaP_ kinetics.

SK channels are more commonly associated with high-threshold Ca^2+^ sources across the CNS ^39, 48, 75–77^, including spinal motor circuits ^19, 78^. Our findings therefore reveal an atypical partnership between SK channels and T-type Ca^2+^ channels, so far reported in midbrain dopaminergic and thalamic neurons ^79, 80^, and extend this mode of coupling to the locomotor CPG. Mechanistically, mild depolarization may activate Cav3.2 channels at subthreshold voltages, producing a local Ca^2+^ signal sufficient to recruit nearby SK conductances. In turn, SK activation would oppose burst emergence by reinforcing AHP-dependent restraint and limiting the depolarizing drive required for regenerative bursting. Consistent with this view, both apamin and T-type antagonists produce similar effects on locomotor output, including burst prolongation and rhythm slowing in lamprey and rodent spinal preparations ^16, 19–23, 72^. Our data extend these observations by showing that local modulation of SK2/3 within the CPG can either initiate (SK inhibition) or terminate (SK potentiation) locomotor rhythms, while T-type Ca^2+^ channel blockade similarly triggers locomotor activity. *In vivo* studies support this bidirectional control; increasing SK activity reduces locomotion in mice ^81^, while SK inhibition in Xenopus embryos increases episode frequency and swimming duration ^73^.

More broadly, SK-T-type coupling may provide a flexible mechanism through which locomotor output is adjusted to behavioral demands. Monoamines illustrate this potential by converging on SK-mediated AHP regulation. Serotonin reduces AHPs in CPG interneurons and motoneurons ^82, 83^, either through direct inhibition of SK channels ^83, 84^ or by limiting Ca^2+^ influx ^85, 86^, thereby prolonging bursts and slowing the rhythm ^87^. Dopamine exerts similar effects ^88, 89^, and noradrenaline may do likewise by reducing Ca^2+^ sensitivity of SK2 gating ^90^. Cholinergic C-bouton inputs onto motoneurons similarly engage muscarinic receptors near SK channels to reduce the AHP and enhance locomotor output ^70, 91, 92^. Beyond neuromodulation, activity-dependent shifts in extracellular ion concentrations may provide an additional layer of control. Local increases in [K^+^]_o_ and decreases in [Ca^2+^]_o_ at locomotor onset ^16^ would reduce the driving forces for both I_KCa_ and I_Ca_, thereby attenuating SK-mediated inhibition and favoring burst emergence. Such ionic changes could therefore provide a rapid gating mechanism for locomotion initiation, complementing the slower neuromodulatory pathways.

In summary, our data identify SK-T-type coupling as a central mechanism controlling the initiation, modulation, and termination of rhythmic bursting in spinal locomotor CPGs. As a tunable brake on burst propensity, SK-T-type coupling defines the operational range of the locomotor network. More broadly, this coupling may represent a conserved biophysical motif for rhythmogenesis, coordinating oscillatory activity across motor systems beyond locomotion.

## Methods

### Animal models

Experiments were performed on Wistar rats and CD-1 IGS mice of either sex. Hb9::eGFP transgenic mice were used for immunohistochemistry and identification of rhythmogenic interneurons. Animals were used at postnatal day (P) 0-2 for fictive locomotion experiments, P5-12 for standard patch-clamp recordings, and P12-20 for AAV-injected mice. Animals were housed under a 12 h light/dark cycle with ad libitum access to water and food at 21-24°C and 40-60% relative humidity. All experimental procedures were conducted in accordance with French regulations (Décret 2010-118) and approved by the local ethics committee (Comité d’Ethique en Experimentation Animale, CEEA-071, authorization No. B1301404 and protocol No. 17485-2018110819197361 and 50133-2024060612594852).

### shRNA construct

An shRNA sequence targeting mouse SK3 (Kcnn3; 5’-TGAGTGACTATGCTCTGATTT-3’) was cloned into an AAV9 vector under control of the U6 promoter, with eGFP expression driven by a CMV promoter for visualization of transduced cells (VectorBuilder, Chicago, IL, USA). A scrambled non-targeting shRNA sequence (5’-CCTAAGGTTAAGTCGCCCTCG-3’) with no homology to known mouse genes served as control. AAV9 particles were produced at titers ≥ 1 × 10^13^ genome copies (GC)/ml and diluted 1:10 in sterile phosphate-buffered saline prior to injection.

### Intrathecal vector delivery

Neonatal mice (P0) were cryoanesthetized and positioned dorsal side up. The intervertebral space was widened by gently flexing the spine. A glass microcapillary preloaded with diluted AAV9 particles was lowered into the center of the T_13_-L_1_ intervertebral space under visual guidance. A total volume of 0.75 µl was slowly injected manually over ∼30 s. Animals recovered on a heating pad and were returned to their home cage. Experiments were performed 12-20 days post-injection to allow adequate transgene expression and knockdown.

### *In vitro* preparations

#### Slice preparation

Lumbar spinal cords were isolated in ice-cold (4 °C) sucrose-based dissection solution containing (in mM): 252 sucrose, 3 KCl, 1.25 NaH_2_PO_4_, 4 MgSO_4_, 0.2 CaCl_2_, 25 NaHCO_3_, 20 D-glucose (pH 7.4, bubbled with 95% O₂/5% CO₂). The spinal cord was embedded in 1% low-melting-point agar, cooled, and mounted on a vibrating microtome (Leica VT1000S, Leica Biosystems, Wetzlar, Germany). Transverse slices (325 µm) were cut from L1-L2 segments and transferred to a holding chamber containing oxygenated (95% O□/5% CO□) aCSF at 30-32 °C composed of (in mM): 120 NaCl, 3 KCl, 1.25 NaH_2_PO_4_, 1.3 MgSO_4_, 1.2 CaCl_2_, 25 NaHCO_3_, 20 D-glucose (pH 7.4). This solution is referred to as standard aCSF ([Ca^2+^]□ = 1.2 mM; [K^+^]□= 3 mM). After 60 min recovery, slices were transferred to a recording chamber continuously perfused with standard aCSF at 32 °C (2-3 ml/min).

#### Whole-spinal cord preparation

The spinal cord was transected at T4, isolated with intact dorsal and ventral roots, and transferred to a recording chamber. The preparation was continuously superfused with oxygenated (95% O₂/5% CO₂) aCSF at 25-26 °C containing (in mM): 120 NaCl, 4 KCl, 1.25 NaH_2_PO_4_, 1.3 MgSO_4_, 1.2 CaCl_2_, 25 NaHCO_3_, 20 D-glucose (pH 7.4).

#### Cell culture

HEK293 cells were maintained in DMEM, high glucose (Gibco) supplemented with 10% fetal bovine serum and penicillin/streptomycin (Gibco), incubated at 37°C in a humidified atmosphere containing 5% CO2. HEK293 cells were transiently transfected using Lipofectamine 3000 (Thermo Fisher) according to the manufacturer’s recommendation. Cells were cotransfected with 2 plasmids: one encoding mKcnn3 (VB241013-1085awr, VectorBuilder) and the other either encoding shRNA targeting mKcnn3 (VB211028-1097hcr) or a control plasmid encoding shRNA against luciferase. The cells were then collected 72 h post transfection.

### *In vitro* recordings

#### Patch-clamp recordings

Whole-cell patch-clamp recordings were performed on ventromedial interneurons in lamina VII-VIII of L_1_-L_2_ segments, adjacent to the central canal, a region enriched in rhythmogenic neurons. Neurons were visualized using infrared differential interference contrast (IR-DIC) videomicroscopy with a 40x water-immersion objective (Olympus) mounted on an upright microscope (Olympus BX51WI) equipped with an infrared-sensitive CCD camera. Hb9::eGFP-positive neurons were identified by epifluorescence. Recordings were performed using a Multiclamp 700B amplifier and Digidata 1550B interface (Molecular Devices). Signals were digitized on-line and filtered at 10 kHz using pClamp 10.7 software (Molecular Devices). Patch pipettes (4-6 MΩ) were pulled from borosilicate glass capillaries (1.5 mm outer diameter, 1.12 mm inner diameter; World Precision Instruments, TW150-4) using a horizontal puller (Sutter P-97). The standard intracellular solution contained (in mM): 140 K^+^-gluconate, 5 NaCl, 2 MgCl_2_, 10 HEPES, 0.5 EGTA, 2 ATP, 0.4 GTP (pH 7.3; 280-290 mOsm). For Ca^2+^ chelation experiments, BAPTA (10 mM) or EGTA (10 mM) replaced the standard 0.5 mM EGTA. In experiments testing the contribution of intracellular Ca^2+^ stores, xestospongin C (2.5 µM) was added to the intracellular solution to block IP_3_ receptors. For experiments requiring K^+^ channel blockade, we use a CsCl-based solution contained (in mM): 120 CsCl, 40 KCl, 2 MgCl2, 10 HEPES, 0.5 EGTA, 2 ATP, 0.4 GTP, pH 7.3. For post hoc morphological identification, biocytin (0.2%) was included in the intracellular solution, and filled neurons were processed as described below (see Immunostaining). The pipette offset potential and capacitance were nulled before seal formation. Data acquisition began at least 5 min after break-in to ensure dialysis and stability. To induce rhythmic activity, the recording chamber was perfused with a modified “locomotor-like” aCSF characterized by reduced calcium and elevated potassium concentrations ([Ca^2+^]□ = 0.9 mM and [K^+^]□ = 6 mM). This ionic environment mimics the extracellular fluctuations observed during the onset of locomotor activity and promotes pacemaker-like membrane oscillations. To block fast synaptic transmission, neurons were isolated from excitatory glutamatergic, glycinergic, and GABAergic inputs using kynurenic acid (1.5 mM) or a combination of CNQX (10 µM) and AP-5 (50 µM), strychnine (1 µM), and picrotoxin (100 µM), respectively. Bicuculline was omitted to avoid blocking the calcium-activated potassium current underlying the afterhyperpolarization ^93^, thereby potentially facilitating burst firing in spinal interneurons ^4^.

#### Extracellular recordings

Fictive locomotion was recorded from L_5_ ventral roots using glass suction electrodes connected to AC-coupled amplifiers (A-M Systems, model 1700). Signals were amplified (1000×), band-pass filtered (70 Hz to 3 kHz), digitized at 5 kHz, and simultaneously rectified and integrated online (t = 100 ms) using custom-built integrators. Locomotor-like activity was induced by bath application of NMDA (5 µM) and 5-HT (10 µM). Prior to pharmacological testing, all spinal cord preparations were first tested for their ability to generate locomotor-like activity in response to NMDA/5-HT. Only preparations showing robust rhythmic ventral root activity were used for subsequent experiments. For focal drug application to the rhythmogenic region, drugs were delivered via pressure ejection through a glass micropipette positioned over the L_1_-L_2_ ventromedial region using a micromanipulator under visual control. The ejection solution contained neutral red to visualize diffusion, and perfusion flow was arranged to prevent drug spread to the ventral root recording sites.

### Immunostaining

Spinal cords from P14-P20 Hb9::eGFP mice were dissected and immersion-fixed for 1 h in 0.25-4% paraformaldehyde (PFA). Tissues were rinsed in phosphate-buffered saline (PBS), cryoprotected overnight in 20% sucrose at 4 °C, and embedded in OCT medium (Tissue-Tek). Transverse cryosections (30 µm) were collected from L_1_-L_2_ segments. Sections were rehydrated in Tris-buffered Saline (TBS) for 15 min at room temperature, permeabilized in TBS containing 10% normal horse serum and 0.4% Triton X-100 for 1 h, and incubated overnight at 4°C in blocking solution (10% normal horse serum, 0.2% Triton X-100 in TBS) with primary antibodies.

Primary antibodies included: polyclonal rabbit anti-SK2 (1:400; #APC-028, Alomone Labs), rabbit anti-SK3 (1:400; #APC-025, Alomone Labs), guinea-pig polyclonal anti-SK2 (1:400; #APC-028-GP, Alomone Labs), and polyclonal rabbit anti-Cav3.2 (1:400; #ACC-025, Alomone Labs). For colocalization assays, sections were co-incubated with guinea-pig polyclonal anti-SK2 and rabbit anti-SK3. Endogenous GFP was enhanced using goat polyclonal anti-GFP (1:400; #ab6673, Abcam).

For biocytin-filled neurons, slices containing recorded cells were fixed immediately after recording in 4% PFA overnight at 4 °C, washed in PBS, and processed as free-floating sections with permeabilization extended to 6 h (0.5% Triton X-100).

After incubation with primary antibodies, sections were washed in TBS and incubated for 1-2 h at room temperature with secondary antibodies: donkey anti-guinea-pig Cy3 (1:400; #706-165-148, Jackson ImmunoResearch), donkey anti-rabbit Cy5 (1:400; #711-175-152, Jackson ImmunoResearch), donkey anti-rabbit Alexa Fluor 555 (1:400; #A31572, Invitrogen), or donkey anti-goat Alexa Fluor 488 (1:400-1:800; #A11055, Life Technologies; #705-545-147, Jackson ImmunoResearch). For biocytin visualization, Alexa Fluor 405-conjugated streptavidin (1:1000; #S32351, Invitrogen) was added. Sections were rinsed, mounted in aqueous mounting medium, and coverslipped. Confocal images were acquired on a Zeiss LSM700 using x20 and x40 oil-immersion objectives. Z-stacks were collected with 0.3-1 µm optical steps and processed in ZEN 12.0 (Zeiss). Laser power, gain, and offset were adjusted for each tissue section to maximize signal-to-noise ratio while avoiding saturation.

### SK3 channel protein quantification

Transfected HEK293 cells were lysed in ice-cold buffer containing 1% Igepal CA-630 and 0.1% SDS, supplemented with protease inhibitors (Complete Mini, Roche Diagnostics). Lysates were centrifuged at 10,000 rpm for 10 min at 4 °C, and the supernatant was used as the total protein fraction. Protein concentrations were determined using a detergent-compatible assay (Bio-Rad). Equal amounts of protein (40 µg per lane) were separated on 4-15% gradient SDS-PAGE stain-free gels (Bio-Rad), transferred to nitrocellulose membranes, and probed overnight at 4 °C with either a polyclonal rabbit anti-SK3 antibody (1:500, Alomone Labs, APC-025) or an anti-actin antibody (1:1,000, A2066, Sigma-Aldrich) in Tris-buffered saline containing 5% fat-free milk. Membranes were then incubated for 1 h at room temperature with a goat anti-rabbit horseradish peroxidase-conjugated secondary antibody (1:40,000; Thermo Fisher).

Immunoreactive bands were visualized using enhanced chemiluminescence detection (Merck-Millipore) and quantified with Image Lab software (Bio-Rad).

### Pharmacological agents

All pharmacological agents were prepared as concentrated stock solutions and diluted to their final concentrations in oxygenated aCSF immediately before use. Compounds were purchased from the following suppliers: Sigma-Aldrich: apamin (100-200 nM), mibefradil (10 µM), NiCl_2_ (200 µM), CdCl_2_ (100 µM), nifedipine (10 µM), veratridine (60 nM), BAPTA (10 mM), EGTA (10 mM), N-methyl-D-aspartate (NMDA, 5 µM), serotonin (5-HT, 10 µM), strychnine (1 µM), picrotoxin (100 µM), biocytin (0.1%), and all salts for aCSF as well as agar/agarose. Alomone: tamapin (5-10 nM), 1-EBIO (200 µM), CyPPA (1-10 µM), and Tram-34 (5 µM). Smartox Biotech: lei-dab7 (5-10 nM), iberiotoxin (200 nM), and ω-Agatoxin IVA (100 nM). Tocris: UCL-1684 (0.2-1 µM), dantrolene (10 µM), xestospongin C (2.5 µM), and riluzole (5 µM). Hello Bio: SNX-482 (100 nM) and kynurenic acid (1.5 mM).

### Computational modeling

#### Hodgkin Huxley Cell Model

For this study we extended a previously described single-compartment Hodgkin-Huxley (HH) type neuronal model ^16, 18^. Membrane potential dynamics were governed by the following equation:

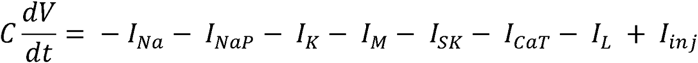

where v is the membrane potential (mV), *l* denotes membrane currents (pA), and *gτ_x_* denotes maximal conductances (nS). Individual ionic currents were defined as:

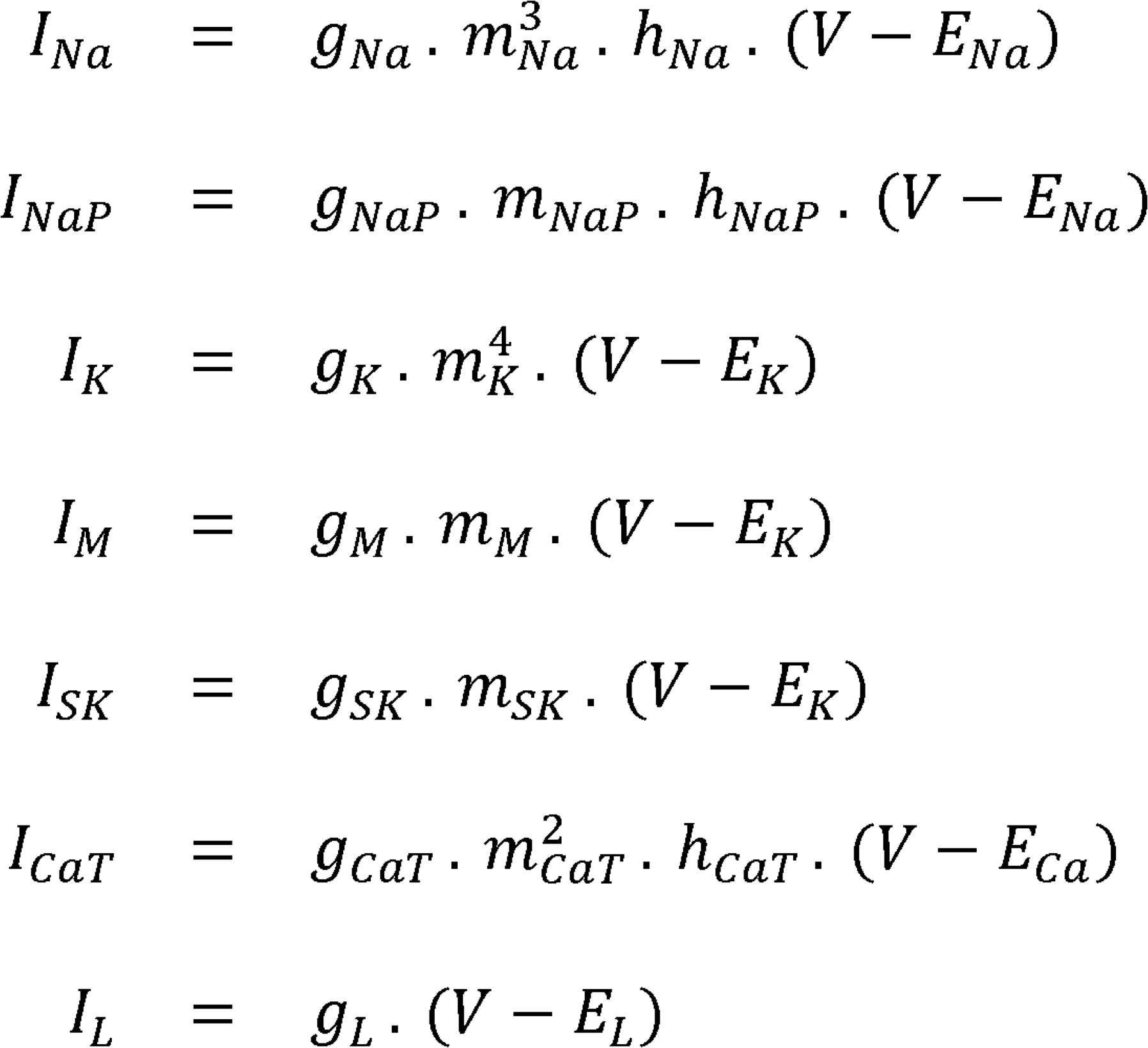

*E_Na_*, *E_K_* and *E_L_* denote the reversal potentials for sodium, potassium, and leak currents, respectively. *E_Na_* and *E_K_* were calculated using the Nernst equation, whereas *E_L_* was computed using the Goldman-Hodgkin-Katz equation based on extracellular (out) and intracellular (in) ionic concentrations. Ionic concentrations were assumed constant throughout the simulation: [*Na*^+^]*_out_* = 145 mM; [*Na*^+^]*_in_* = 15 mM; [*K*^+^]*_out_* = 3mM;[*K*^+^]*_in_* = 140 mM; [*Cl^-^*]*_out_* = 130 mM; [*Cl^-^*]*_in_* = 8mM. Relative permeabilities of sodium and chloride ions (with respect to potassium ions) were set to 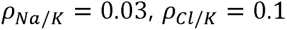.

Activation and inactivation variables followed standard first-order kinetics:

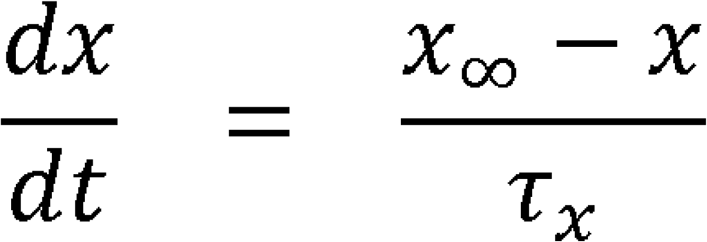

Where 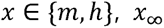 represents the voltage-dependent steady-state value, and *τ_x_* is the voltage-dependent time constant. These quantities are defined as:

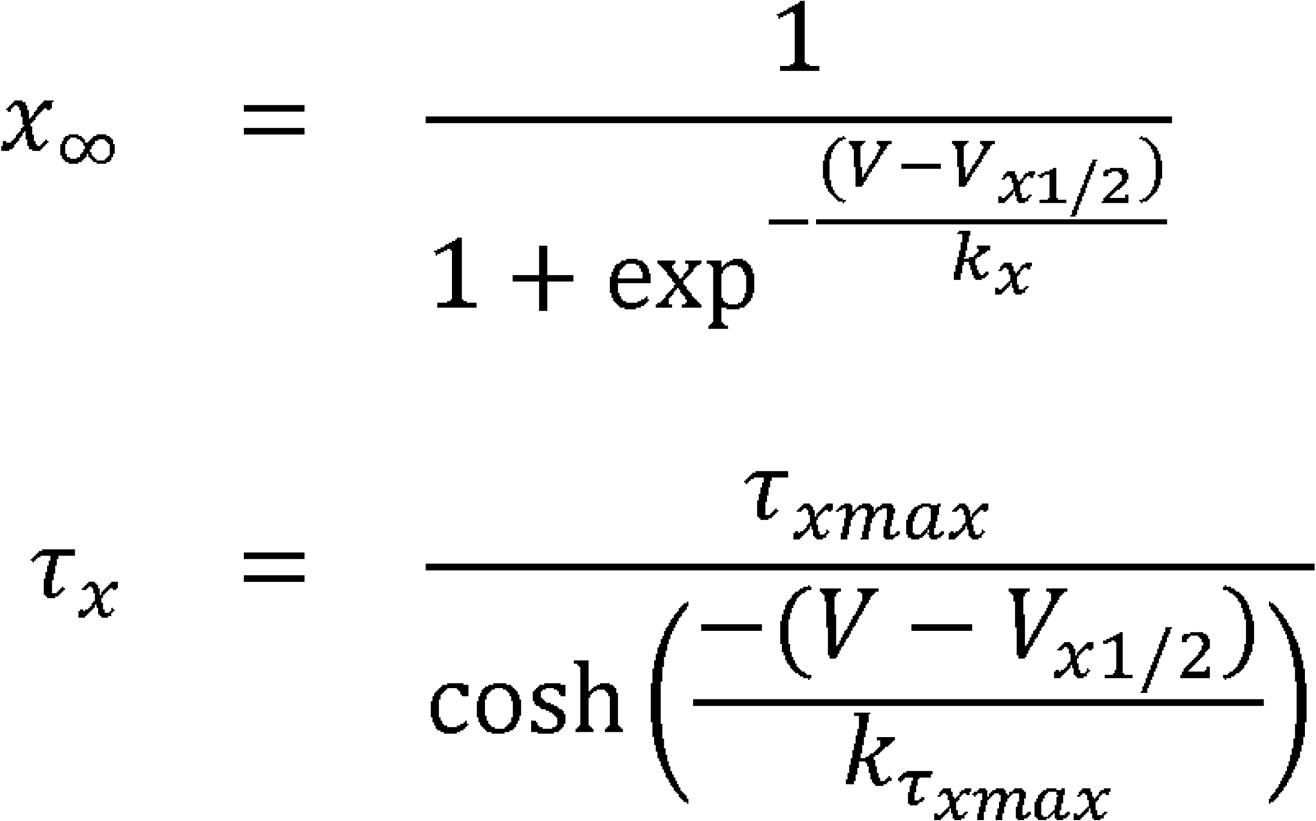

where, *V_x1/2_* and *k_x_* denote the half-activation voltage and the slope factor, respectively, and *τ_xmax_* and *kτ* define the voltage dependence of the corresponding time constant. Activation kinetics for *l_Na_* and *l_NaP_* were assumed instantaneous 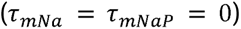, such that *m_Na_* and *m_NaP_* were set to steady-state values.

Unlike the voltage-dependent channels above, SK-channel gating depended on intracellular calcium concentration rather than membrane voltage. Intracellular calcium dynamics and SK activation were described as:

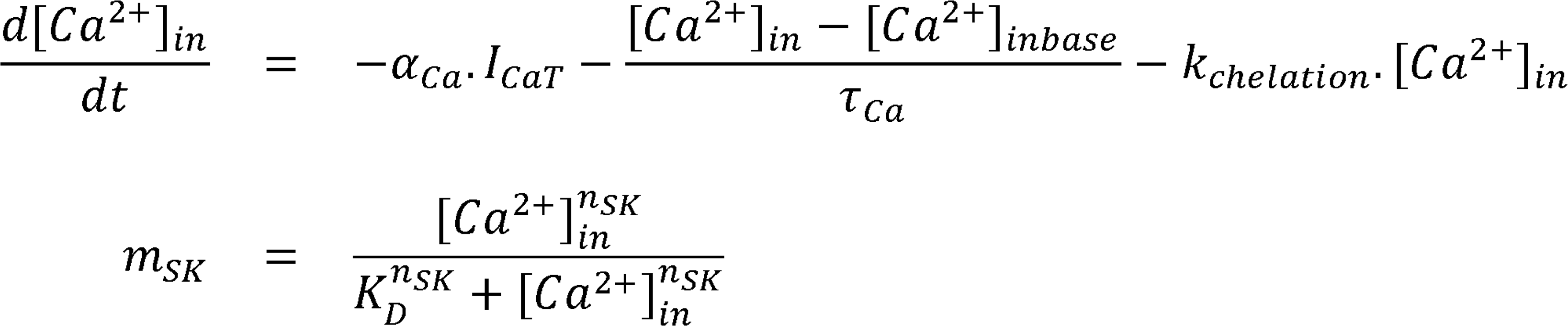

Because SK-channel activation is faster than intracellular calcium dynamics, it was assumed to be instantaneous. Here *α_ca_* converts calcium current *I_ca_* (pA) into intracellular concentration change 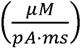. Extracellular calcium concentration was set to [*Ca^2^*^+^]*_out_* = 1.2 nM, basal intracellular calcium concentration to [*Ca^2^*^+^]*_inbase =_* 50 nM, and the calcium reversal potential fixed to *E_ca_* = 120 mV.

To capture trial-to-trial variability, the injected current included a temporally correlated noise term modeled as an Ornstein-Uhlenbeck process, adapted from ^94^.

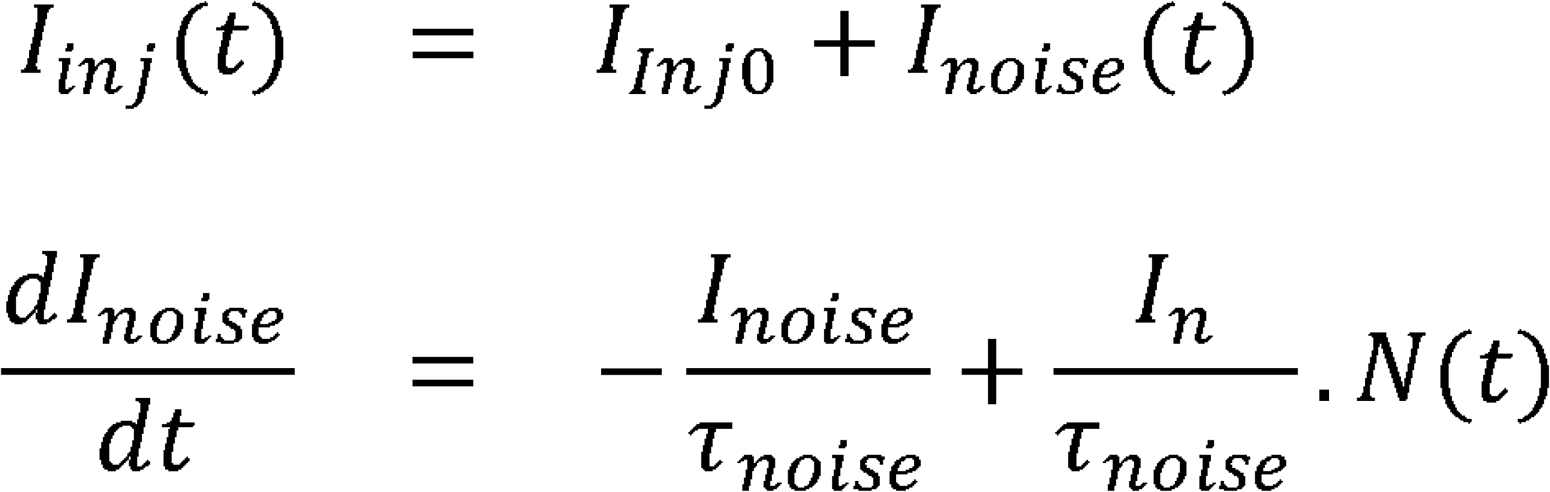

where *N*(*t*) is a normally distributed random variable 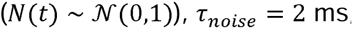, and *l_n_* = 5. All model parameters are summarized in Table 1.

**Table 1.**
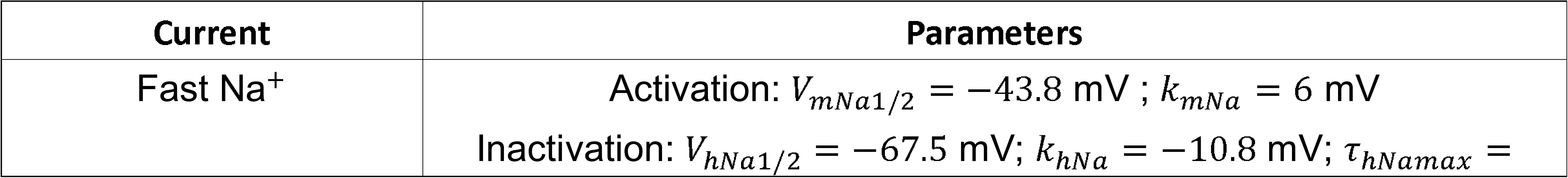
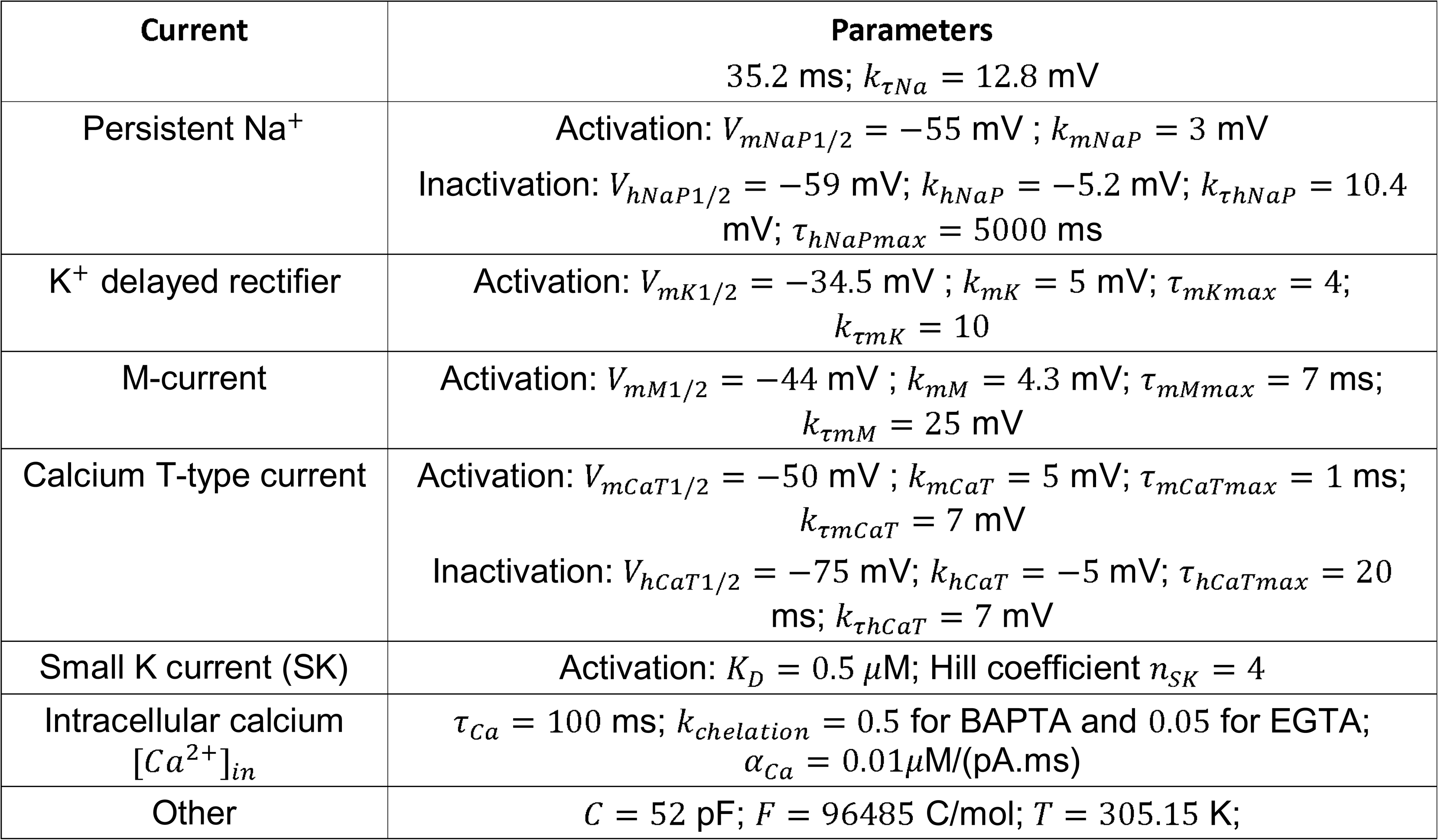
Biophysical parameters of the extended Hodgkin–Huxley model.

#### Simulation-Based Inference (SBI)

To fit the biophysical model to experimental data while accounting for parameter uncertainty, we used simulation-based inference (SBI), a Bayesian framework for parameter estimation ^95^. SBI estimates posterior distributions over model parameters from observed summary features extracted from neuronal activity. Here, the inferred parameters initially comprised the six conductances of the Hodgkin-Huxley model. Ten summary features were computed from the voltage envelope of each burst phenotype: burst duration, burst duration variance, maximum burst amplitude, inter-burst frequency, area under the curve (AUC), interburst period, the maximum and minimum of the first derivative of the envelope, the maximum and minimum of the envelope acceleration, and the minimum and maximum membrane depolarization.

Unlike classical Bayesian approaches, SBI does not require explicit evaluation of the likelihood function relating model parameters to observed features, which is generally intractable for nonlinear systems such as the present Hodgkin-Huxley model. SBI circumvents this by approximating the likelihood using a normalizing flow, a class of deep generative models trained on forward simulations to learn an invertible mapping between parameter and feature spaces ^96^. To this end, parameter sets were sampled from prior distributions defined over physiologically plausible ranges for the six conductances. Priors were uniform and identical across the three clusters. In total, 20,000 parameter configurations were simulated, and the corresponding voltage traces were generated using a Runge-Kutta integration scheme. Numerical simulation was performed using Runge-Kutta integration method for stochastic processes, as the injected current to the neuron (I_inj_) involved randomness, chosen to be of constant amplitude.

Inference quality was assessed using standard posterior diagnostics. Posterior shrinkage, defined as the reduction in parameter uncertainty from prior to posterior, was used to evaluate how strongly individual parameters were constrained by the observed features. Posterior predictive checks further verified that simulations drawn from the inferred posteriors reproduced the target features within acceptable margins. Pairwise posterior distributions were additionally examined to assess parameter dependencies and residual identifiability issues.

For the burst phenotype characterized by sustained plateau depolarization (Cluster 3; see Results), the initial six-parameter inference did not reproduce the waveform satisfactorily. We therefore introduced the maximal time constant governing deinactivation of the I_NaP_ inactivation gate (τ*_hNaPmax_*; see Hodgkin-Huxley model section) as a seventh free parameter, motivated by previous experimental evidence that changes in I_NaP_ kinetics strongly affect burst duration ^6^. This parameter was sampled from a uniform prior over 4,000-80,000 ms, whereas all other priors were kept unchanged.

### *In silico* perturbation analyses

Inferred parameter sets for each cluster were used to perform targeted *in silico* perturbations. In order to reproduce experimental results four perturbation paradigms were examined. First, SK conductance (*g_SK_*) was progressively reintroduced to test whether restoration of SK current suppresses bursting and promotes tonic spiking. Second, persistent sodium conductance (*g_NaP_*) was set to zero to assess whether I_NaP_ is required for burst generation. Third, starting from the SK-restored condition in which bursting had been suppressed, T-type calcium conductance (*g_Caτ_*) was removed to test whether T-type calcium channel blockade was sufficient to restore bursting. Fourth, again from the SK-restored tonic-spiking condition, intracellular calcium handling was perturbed by modifying the chelation parameter *k_chelation_* to mimic the effects of fast and slow calcium chelators. Because the fitted parameter regimes differed across clusters, *in silico* perturbations were implemented using cluster-specific parameter values, summarized in Table 2.

**Table 2.** Cluster-specific parameter values used for in silico perturbation analyses.

| Experiment | Cluster 1 | Cluster 2 | Cluster 3 |
| --- | --- | --- | --- |
| SK reactivation | $g_{SK} = 3.5$ | $g_{SK} = 3.5; I_{inj} = 28$ | $g_{SK} = 3.5; I_{inj} = 22$ |
| $I_{Nap}$ suppression | $g_{NaP} = 0, I_{inj} = 31.5$ | $g_{NaP} = 0; I_{inj} = 33$ | $g_{NaP} = 0; I_{inj} = 26$ |
| CaT suppression | $g_{SK} = 3.5; g_{CaT} = 0$ | $g_{SK} = 3.5; g_{CaT} = 0$ | $g_{SK} = 3.5; g_{CaT} = 0$ |
| Calcium chelation | $g_{SK} = 3.5; k_{chelation} = 0.05$ | $g_{SK} = 3.5; k_{chelation} = 0.05$ | $g_{SK} = 3.5; k_{chelation} = 0.05$ |

### Quantification and statistical analysis

#### Data analysis

Electrophysiological data were analyzed off-line. For whole-cell recordings, several basic criteria were set to ensure optimum quality of intracellular recordings. Only cells exhibiting a stable resting membrane potential and an action potential amplitude larger than 40 mV were considered. Passive membrane properties of cells were measured by determining from the holding potential the largest voltage deflections induced by small current pulses that avoided activation of voltage-sensitive currents. We determined input resistance by the slope of linear fits to voltage responses evoked by small positive and negative current injections. Firing properties were measured from depolarizing current pulses of varying amplitudes. Action potential (AP) features were extracted with a custom Python pipeline (PyABF/SciPy). For each cell, the first spike at rheobase was analyzed. AP threshold was defined as the first sample within the 4 ms preceding the AP peak where dV/dt ≥ 10 mV·ms^-1^. AP amplitude was measured from threshold to peak. AP half-width was computed at half-amplitude with linear interpolation at the rising and falling crossings. Depolarizing and repolarizing slopes (mV·ms^-1^) were computed from threshold to peak and from peak to half-amplitude return, respectively. Instantaneous firing frequency was the reciprocal of interspike interval; a mean instantaneous frequency was calculated across spikes in each sweep. The medium afterhyperpolarization (mAHP) amplitude was measured as the voltage drop from threshold to the most negative membrane potential occurring after the spike. mAHP duration was defined as the time the membrane potential remained below the half-recovery level (midpoint between the mAHP minimum and the threshold voltage), with crossings estimated by interpolation. All reported membrane potentials were offline-corrected for the liquid junction potential, computed for each experiment from the ionic composition of the intra- and extracellular solutions.

Burst features were quantified from bursts evoked at rheobase current. Bursts were identified on raw membrane potential recordings as discrete episodes of action potential firing (active) separated by spike-free (silent) intervals. In some cells, spikes occurred only at burst onset and were followed by a depolarization block during the plateau; these events were still classified as bursts. Bursting properties were then quantified from low-pass filtered (3 Hz) and median-smoothed recordings to remove spikes while preserving the depolarizing envelope. Burst amplitude and duration were measured using a threshold-based detection algorithm, which identified the burst onset, peak, and offset, as well as the corresponding maximum and minimum values of the derivative (dV/dt). Onset and offset were refined by tracking the sign of dV/dt around the threshold crossings. For each burst, amplitude was measured as the maximal depolarization above the onset baseline; duration was the interval from refined onset to offset; area was the Simpson-integrated baseline-subtracted envelope between onset and offset. Burst frequency was calculated as the inverse of the interval between consecutive burst onsets. Quantitative burst features (duration, amplitude, frequency, and maximal depolarizing and hyperpolarizing slopes) extracted from individual neurons were subjected to principal component analysis (PCA) to identify dominant axes of variability. The first two components captured most of the variance and were used for visualization and interpretation. Hierarchical agglomerative clustering using Ward’s linkage was then applied to the standardized variables, revealing three distinct bursting phenotypes. These analyses were performed in Python using scikit-learn and SciPy, and cluster assignments were projected onto the PCA space.

For extracellular recordings, alternating activity between right/left L5 recordings was taken to be indicative of fictive locomotion.

Quantitative image analysis was performed in FIJI (ImageJ, NIH). For somatic analysis, ROIs encompassing the somata of Hb9::eGFP-positive neurons or unidentified ventromedial interneurons located in the L₁-L₂ region near the central canal were manually delineated. For dendritic analysis, ROIs were drawn along the full extent of dendrites from biocytin-filled Hb9::eGFP-positive neurons. Within each ROI, SK2-, SK3-, and Cav3.2-immunopositive puncta were segmented using automated local thresholding after background subtraction. Puncta were then detected using the Analyze Particles plugin with compartment-specific parameters (40× objective). For the quantification of somatic clusters, inclusion criteria were adapted to the target protein. For SK channels, particles were included if they fell within a size range of 0.2-5 µm^2^ with a circularity between 0.75 and 1.0. For Cav3.2 channels, detection parameters were adjusted to a size range of 0.04-0.5 µm^2^ with a circularity between 0.55 and 1.0. For dendritic SK clusters, filtering criteria were set to a size range of 0.12–2 µm^2^ and a circularity of 0.55-1.0. Puncta density and mean particle diameter were extracted from the Analyze Particles outputs. Colocalization was assessed by generating binary masks for each channel and computing their intersection (logical AND) in ImageJ, which isolates overlapping puncta. The intersection mask was then analyzed with Analyze Particles using the same compartment-specific criteria.

### Statistics

Statistical analyses were performed using GraphPad Prism 7 and Python (with the statsmodels and scikit-learn libraries). For two-group comparisons, Wilcoxon, Mann-Whitney or Kolmogorov-Smirnov tests were used as appropriate. The analysis of the estimated posteriors conductances distributions was conducted using the Kolmogorov-

Smirnov test with 50 randomly drawn samples. Multiple groups were analyzed using Kruskal-Wallis or Friedman tests with Dunn’s post hoc correction, and proportions were compared using Fisher’s exact test. One-way ANOVA was applied when data met parametric assumptions. Multivariate analysis of variance (MANOVA) was used to assess the effect of experimental factors on multiple dependent variables (PC1 and PC2 scores) simultaneously. Hierarchical agglomerative clustering was applied to the PCA-transformed dataset, using Ward’s linkage to ensure internal cluster homogeneity. All tests were two-sided, with *P* < 0.05 considered significant. Specific statistical tests, sample sizes (*n*), and *P* values for each dataset are provided in the corresponding figure legends.

## Data availability

All relevant data supporting the findings of this study are included in the Source Data or available from the corresponding authors upon request. Further information and requests for resources and reagents should be directed to and will be fulfilled by the corresponding author, Frédéric Brocard.

## Acknowledgements

We are grateful to Geneviève Rougon for her valuable input to the manuscript. We thank the GeneXprINT platform at the Institut de Neurosciences de la Timone for genotyping services, Anne Duhoux and Mélanie Huc for excellent animal care. This research was supported by Agence National de la Recherche Scientifique (SpasT-SCI-T ANR-21-CE17-0060; MotoBIS ANR-24-CE16-1548 and RhythMIC ANR-24-CE14-4160 to F.B.).

## Author Contributions

F.K. designed, performed and analyzed the vast majority of *in vitro* electrophysiological experiments, contributed to computational modeling and wrote the first draft of the manuscript. C.D. designed, performed and analyzed computational modeling experiments and simulation-based inference analyses under the supervision of M.G. C.B. designed, performed and analyzed immunohistochemistry experiments related to SK2 and SK3 channels. V.T. designed, performed and analyzed immunohistochemistry experiments related to Cav3.2 and validated the shRNA knockdown in cell culture. B.D. performed some intracellular recordings. J.-D.L. and M.H. contributed to the development of the model fitting pipeline.

M.G. designed, performed, analyzed and supervised computational modeling experiments. F.B. conceptualized, administrated, designed, supervised and funded the whole project, updated the Hodgkin-Huxley model, performed and analyzed some *in vitro* experiments and wrote the manuscript.

