## Supplemental Information for "SK2/3 CHANNELS COUPLE WITH T-TYPE CA^2+^ CHANNELS TO GATE SPINAL LOCOMOTOR RHYTHM GENERATION"

**SUPPLEMENTARY INFORMATION**

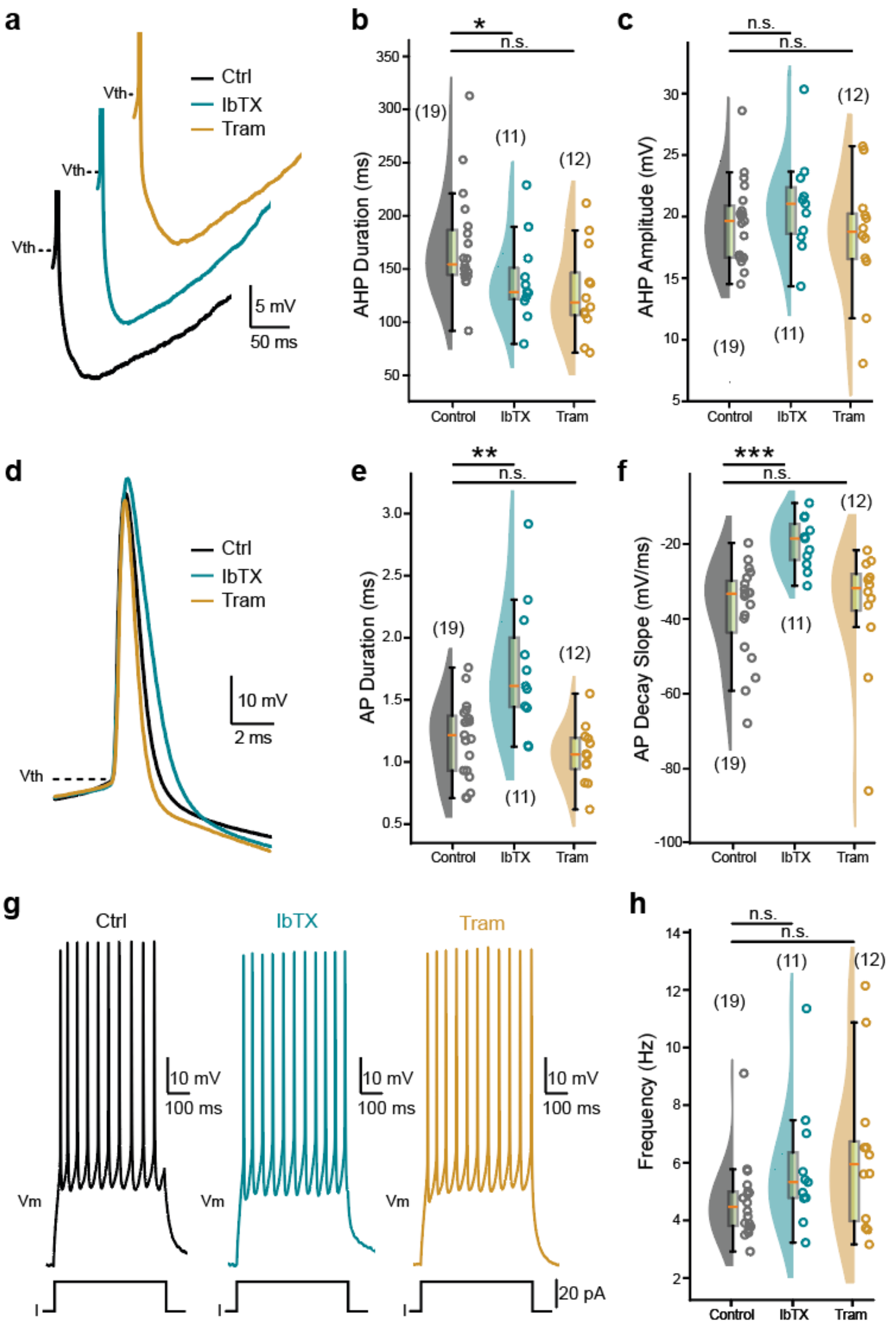


**Supplementary Figure 1. BK and IK channels exhibit no role in burst induction.** **a, d, g** Representative voltage traces from ventromedial interneurons ($L_1$-$L_2$) evoked by depolarizing pulses under control conditions (black), or in the presence of IbTx (teal) or Tram-34 (gold). Panels display individual afterhyperpolarizations (AHP; **a**), single action potentials (**d**) and repetitive spiking (**g**). Dashed lines indicate the spiking threshold ($V_{Th}$). **b, c, e, f, h** Raincloud plots with box-and-whisker overlays (median, interquartile range) comparing AHP half-width duration (**b**) and amplitude (**c**), action potential duration (**e**) and decay slope (**f**) and firing frequency (**h**) across control, IbTx, and Tram-34 conditions. Numbers in parentheses denote recorded cells; each dot a single cell. n.s., not significant; ***P* < 0.01; ****P* < 0.001 (Kruskal-Wallis with Dunn's post hoc test for **b, c, e, f, h**). For detailed *P* values, see Source data.

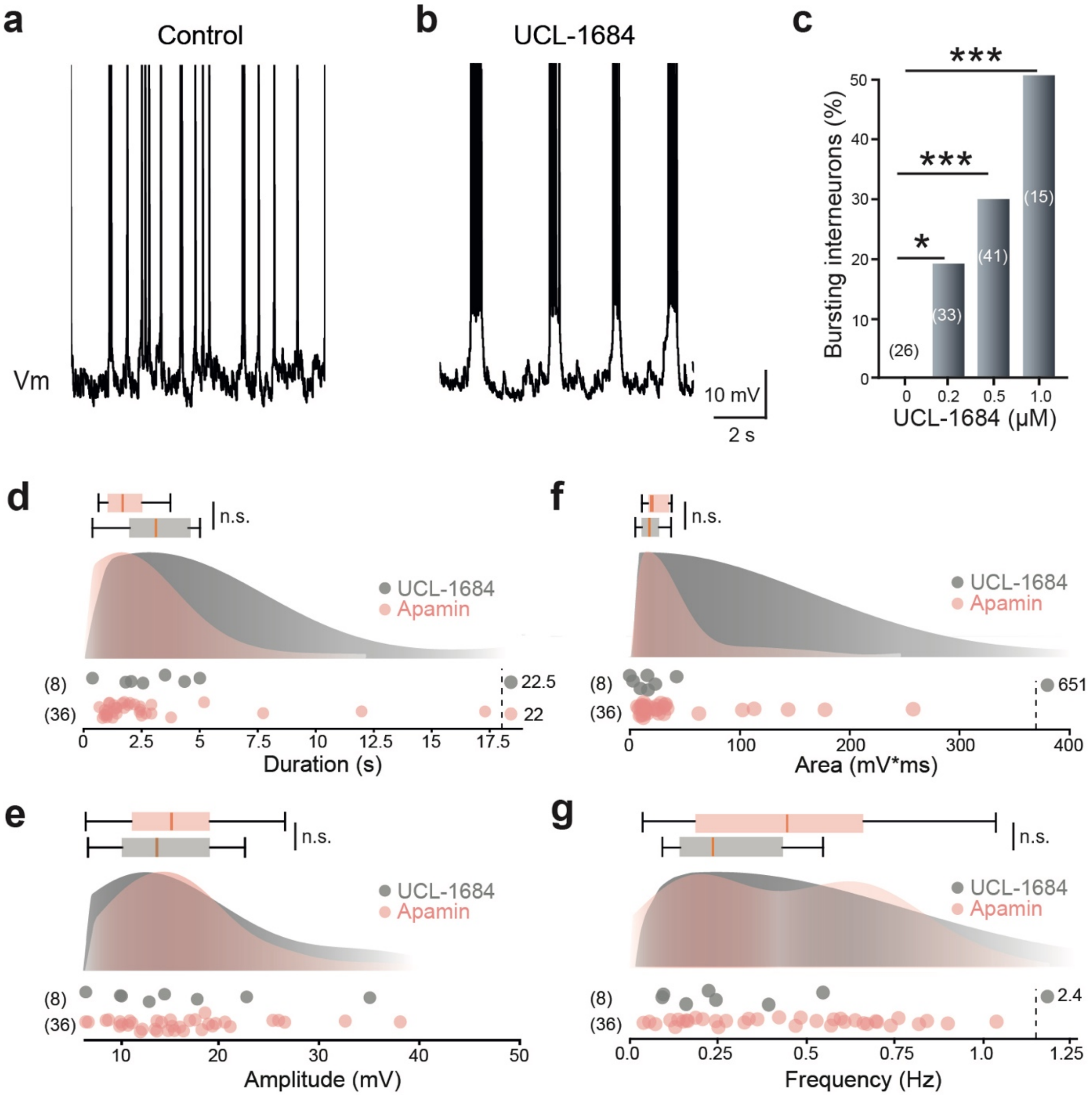


**Supplementary Figure 2. The broad-spectrum SK channel blocker UCL-1684 induces bursting.** **a, b** Voltage traces from ventromedial interneurons ($L_1$-$L_2$) under control conditions (**a**) and after bath application of UCL-1684 (1 μM, **b**). **c** Bar graph showing the percentage of bursting interneurons recorded at increasing concentrations of UCL-1684 (0.2-1 μM). **d-g** Raincloud plots with box-and-whisker overlays (median, interquartile range) comparing burst duration (**d**), amplitude (**e**), area (**f**), and frequency (**g**) recorded under UCL-1684 (1 μM; grey) or apamin (pink). Data points plotted beyond dashed vertical lines indicate values outside the axis range. Numbers in parentheses denote recorded cells; each dot represents a single cell. n.s., not significant; *$P$ < 0.05; **$P$ < 0.001 (two-sided Fisher's exact test for **c**; two-sided Mann-Whitney test for **d-g**). For detailed $P$ values, see Source data.

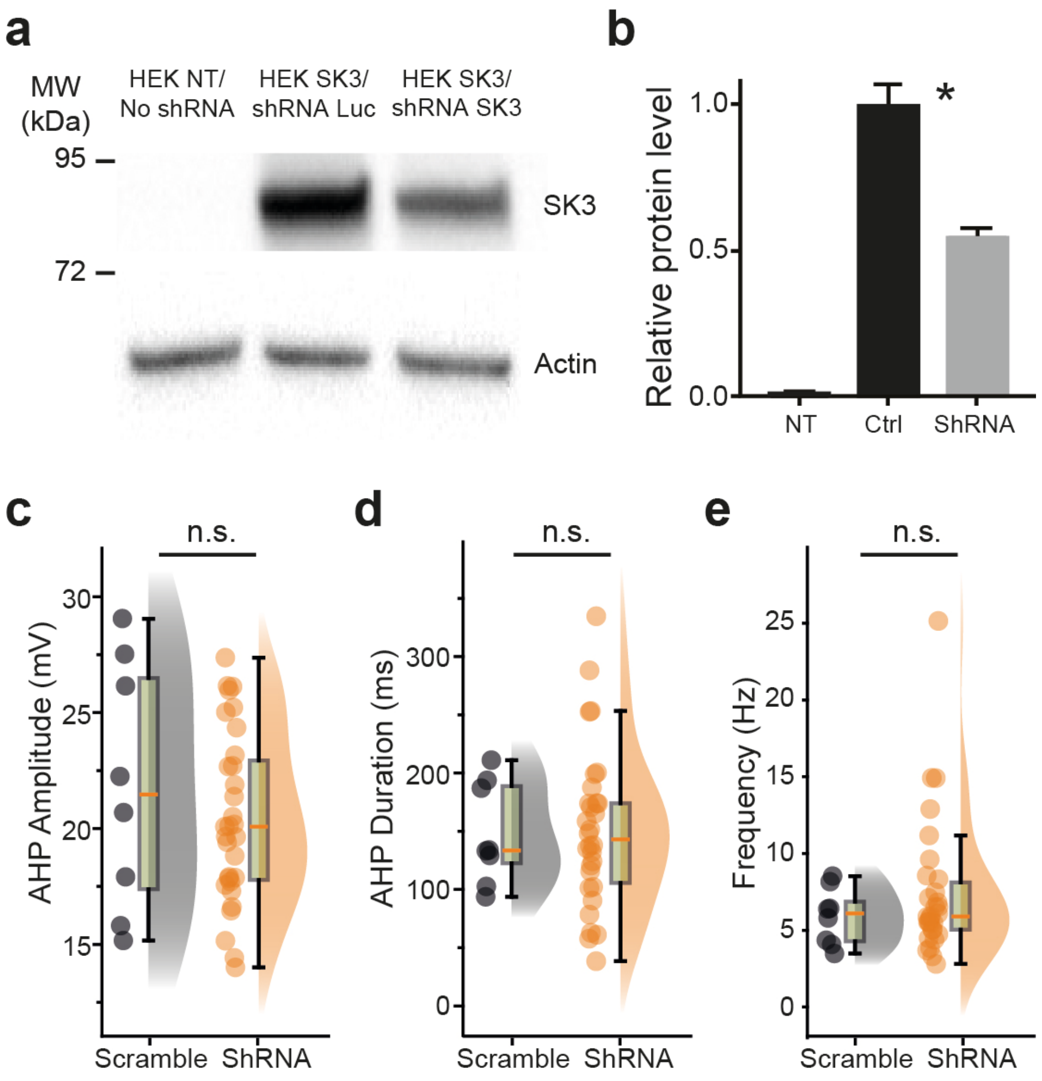


**Supplementary Figure 3. Validation of SK3 knockdown efficiency.** **a** Western blot showing SK3 protein expression (upper panel) and actin loading control (lower panel) in non-transfected cells (NT), cells co-expressing SK3 and control shRNA (shLuc), and cells co-expressing SK3 and SK3-targeting shRNA. **b** Bar graph quantifying SK3 protein levels normalized to actin and expressed relative to control shRNA (n = 4 biological replicates). **c-e** Raincloud plots with box-and-whisker overlays (median, interquartile range) comparing AHP amplitude (**c**), AHP duration (**d**), and firing frequency (**e**) in interneurons transfected with control shRNA (scramble; grey) or SK3-targeting shRNA (orange). Note that unlike SK2 blockade, SK3 knockdown does not alter AHP properties. Numbers in parentheses denote recorded cells; each dot represents a single cell. n.s., not significant; * $P < 0.05$. (two-sided Mann-Whitney test for **b-e**). For detailed P values, see Source data.

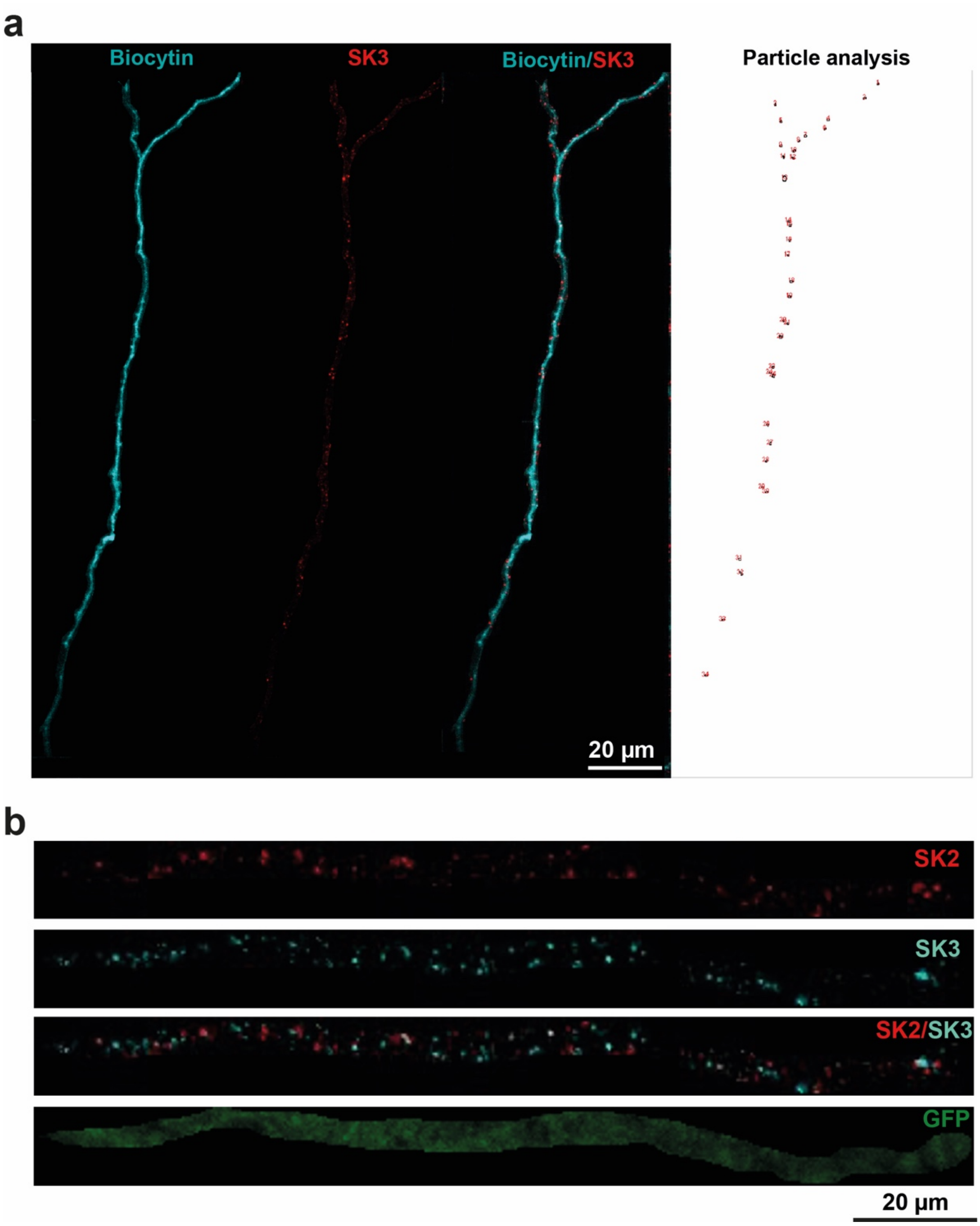


**Supplementary Figure 4. Dendritic SK channel cluster analysis in Hb9 interneurons.** **a** Representative confocal images of a dendritic branch from an intracellularly recorded Hb9 interneuron filled with biocytin (cyan), showing SK3 immunolabeling (red), the merged overlay, and the corresponding particle analysis (red dots) used for cluster quantification. Scale bar, 20 µm. **b** High-magnification confocal images of a dendritic segment from an Hb9-GFP interneuron, showing SK2 immunolabeling (red), SK3 immunolabeling (cyan), the merged SK2/SK3 signal, and GFP fluorescence (green). Scale bar, 20 µm.

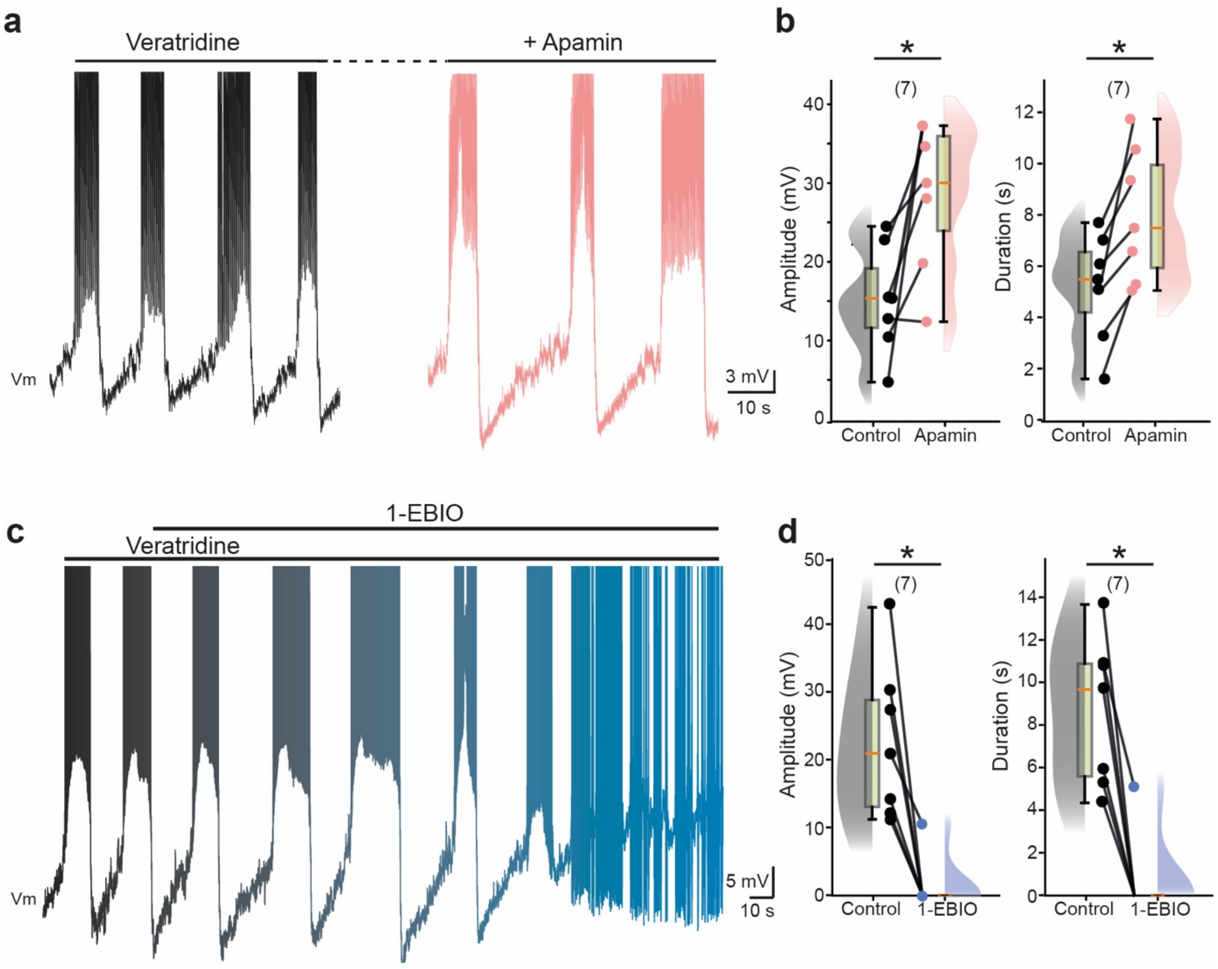


**Supplementary Figure 5. SK activation and blockade oppositely regulate veratridine-induced $I_{NaP}$-dependent bursting.** **a, c** Voltage traces from ventromedial interneurons of the rhythmogenic CPG region ($L_1$-$L_2$) recorded in the presence of veratridine (60 nM; black) before and after bath application of the SK channel blocker apamin (200 nM; pink, **a**) or the SK activator 1-EBIO (200 µM; blue, **c**). **b, d** Raincloud plots with box-and-whisker overlays (median, interquartile range) quantifying burst amplitude and duration under veratridine control conditions versus veratridine co-applied with apamin (**b**) or 1-EBIO (**d**). Numbers in parentheses denote recorded cells; each dot represents a single cell. **P* < 0.05 (two-sided Wilcoxon paired test). For detailed P values, see Source data.

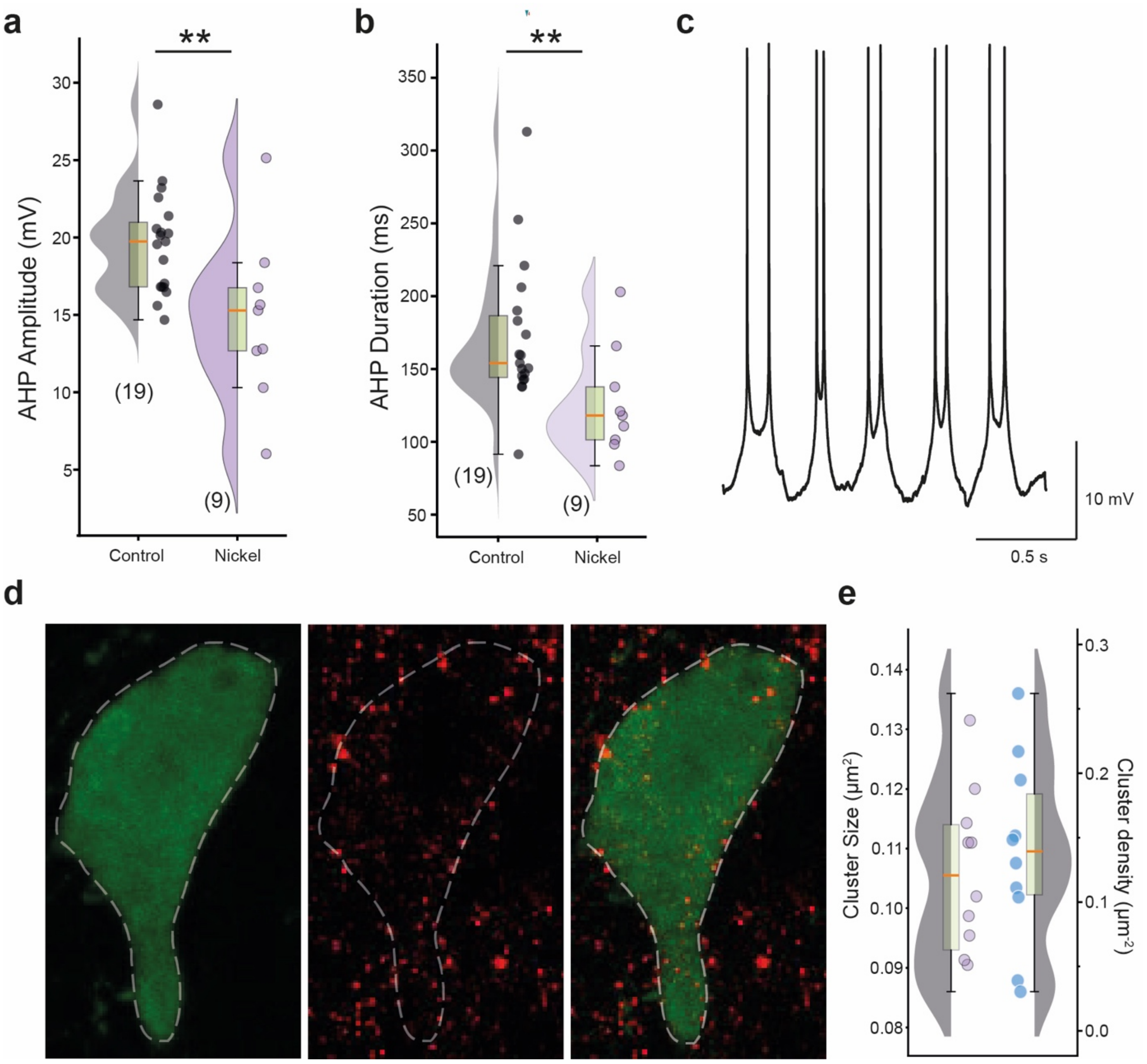


**Supplementary Figure 6. The highly nickel-sensitive Cav3.2 channels contribute to burst dynamics in Hb9 interneurons.** **a**, **b**, Raincloud plots with box-and-whisker overlays (median and interquartile range) showing AHP amplitude (**a**) and AHP duration (**b**) under control conditions (grey) and after nickel application (100 µM, purple). **c**, Representative voltage trace illustrating rhythmic bursting recorded in the presence of $Ni^{2+}$ (100 µM). **d**, Confocal images of an Hb9-GFP interneuron showing GFP fluorescence (*left*), Cav3.2 immunolabeling (*middle*), and merged overlay (*right*). Cav3.2 immunoreactivity forms discrete clusters along the somatic membrane (dashed line). Scale bar, 5 µm. **e**, Raincloud plots showing Cav3.2 cluster size (left axis, purple dots) and cluster density (right axis, blue dots) measured on the somatic membrane of Hb9-GFP interneurons. Numbers in parentheses denote recorded cells (**a, b**) or analyzed neurons (**e**); each dot represents a single cell. ** $P < 0.01$ (two-sided Mann-Whitney test for **a, b**). For detailed P values, see Source data.

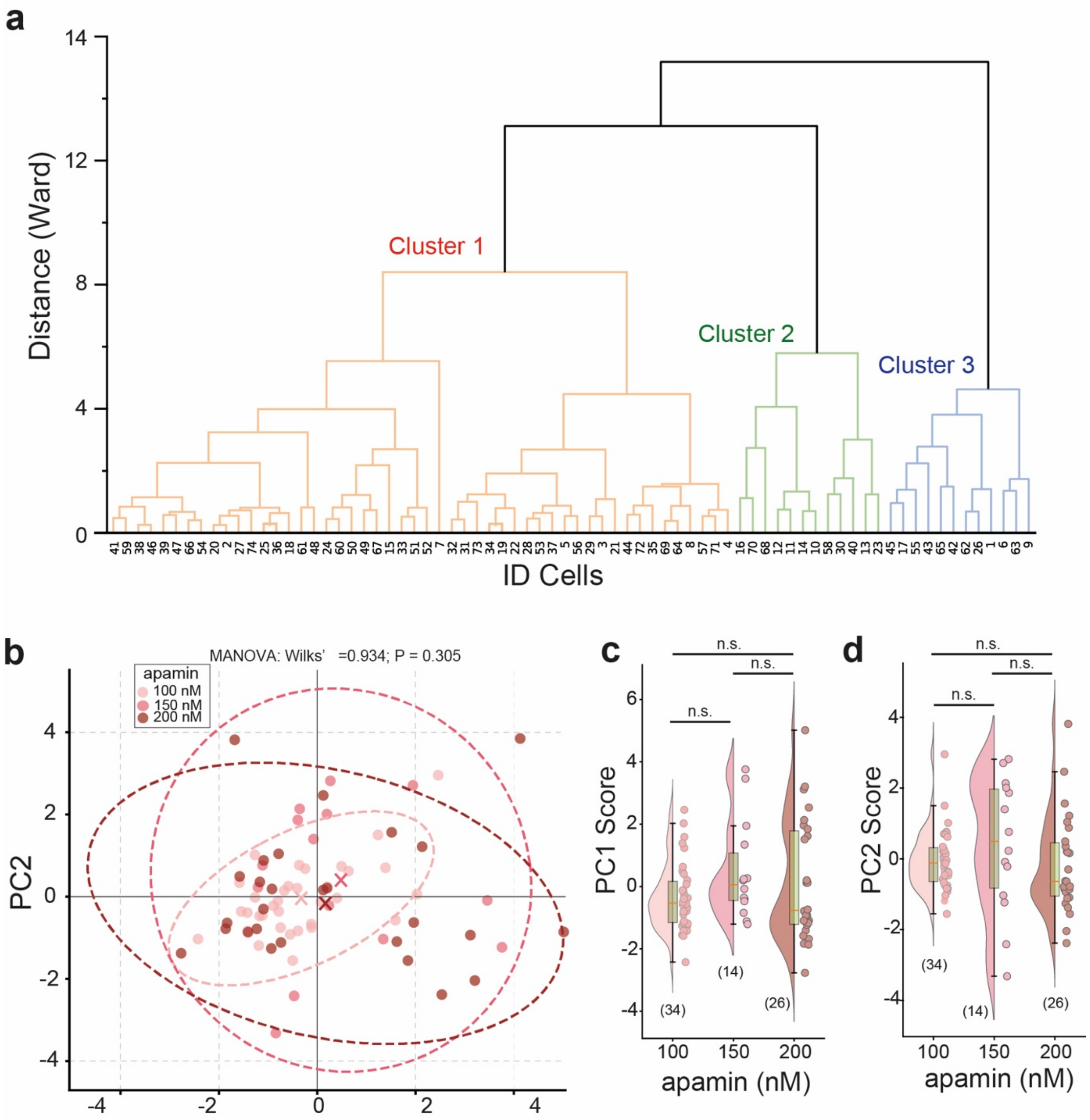


**Supplementary Figure 7. Burst phenotypes are independent of apamin concentration. a** Hierarchical clustering dendrogram (Ward linkage) computed from burst features, with branches colored according to the three clusters (Cluster 1, orange; Cluster 2, green; Cluster 3, blue). **b** Principal Component Analysis (PCA) projection of burst characteristics (PC1 vs. PC2) grouped by apamin concentration (100, 150, or 200 nM). Crosses indicate group centroids (multivariate means), and dashed ellipses represent 95% confidence intervals. Note the substantial overlap between groups, indicating no significant dose effect (MANOVA, Wilks' lambda, P = 0.305). **c, d** Raincloud plots with box-and-whisker overlays (median, interquartile range) comparing PC1 (**c**) and PC2 (**d**) scores across the three apamin concentrations. Numbers in parentheses denote recorded cells; each dot represents a single cell. n.s., not significant (multivariate analysis of variance (MANOVA with Wilks' lambda for **b**, and one-way ANOVA for **c**). For detailed P values, see Source data.

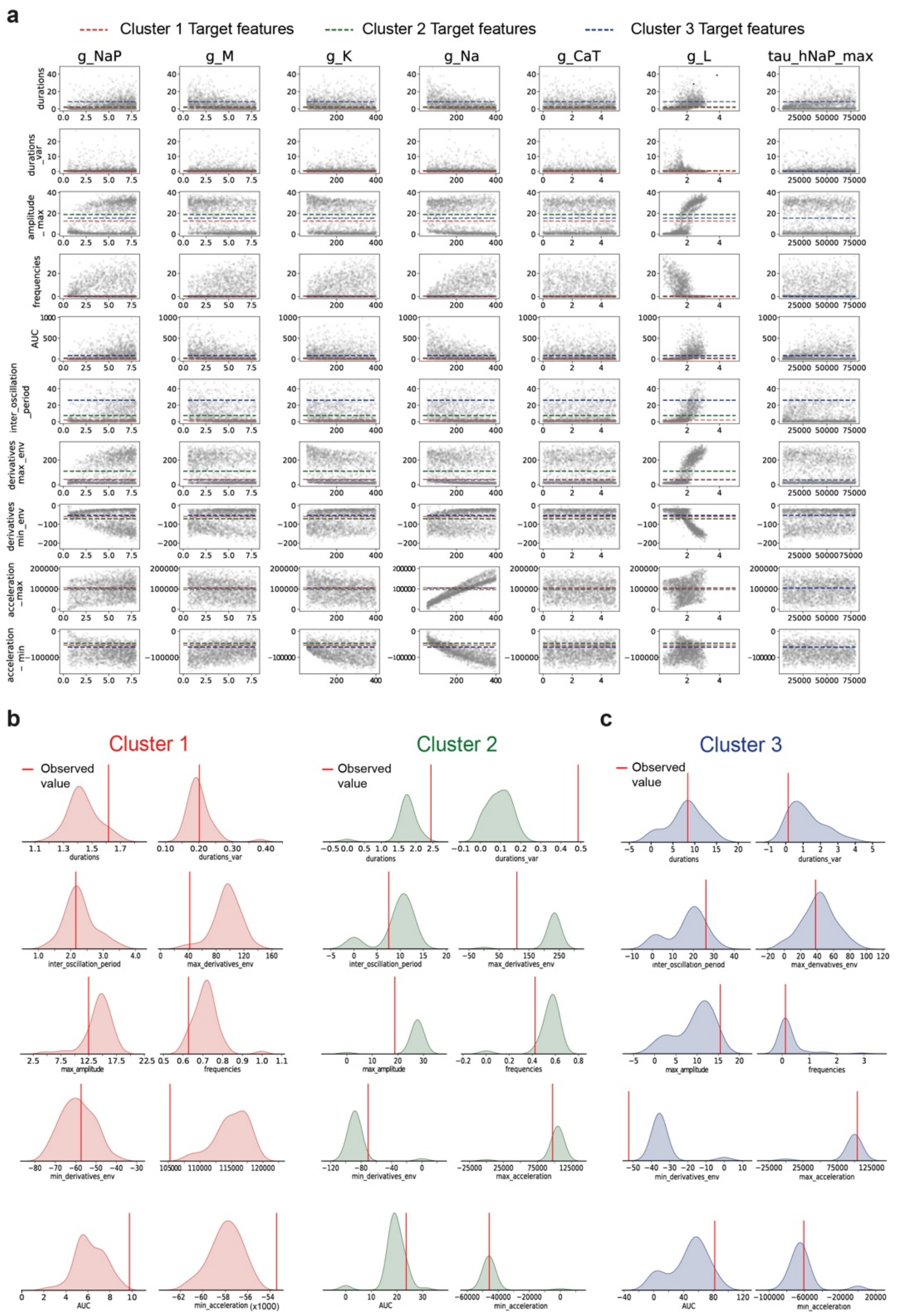


**Supplementary Figure 8. Prior and posterior predictive checks for simulation-based inference. a** Prior predictive check. Scatter plots show the values of each summary feature (y-axis, one feature per row) as a function of each model parameter (x-axis, one parameter per column) for 2,000 representative simulations subsampled from 20,000 total. Colored dashed lines indicate the target feature values for each cluster (Cluster 1, red; Cluster 2, green; Cluster 3, blue). The rightmost column shows the sensitivity of features to the $I_{NaP}$ deinactivation time constant ($\tau_{hNaPmax}$), included as an additional free parameter to capture the long burst durations of Cluster 3. **b** Posterior predictive checks for Cluster 1 (red) and Cluster 2 (green). Distributions represent feature values obtained from 100 simulations drawn from the initial six-parameter posterior distributions. **c** Posterior predictive checks for Cluster 3 (blue). Because the initial six-parameter inference failed to satisfactorily reproduce the sustained plateau depolarization of Cluster 3, $\tau_{hNaPmax}$ was introduced as a seventh free parameter. Distributions show feature values obtained from 100 simulations drawn from this expanded seven-parameter posterior distribution. Red vertical lines indicate the observed target value for each feature.

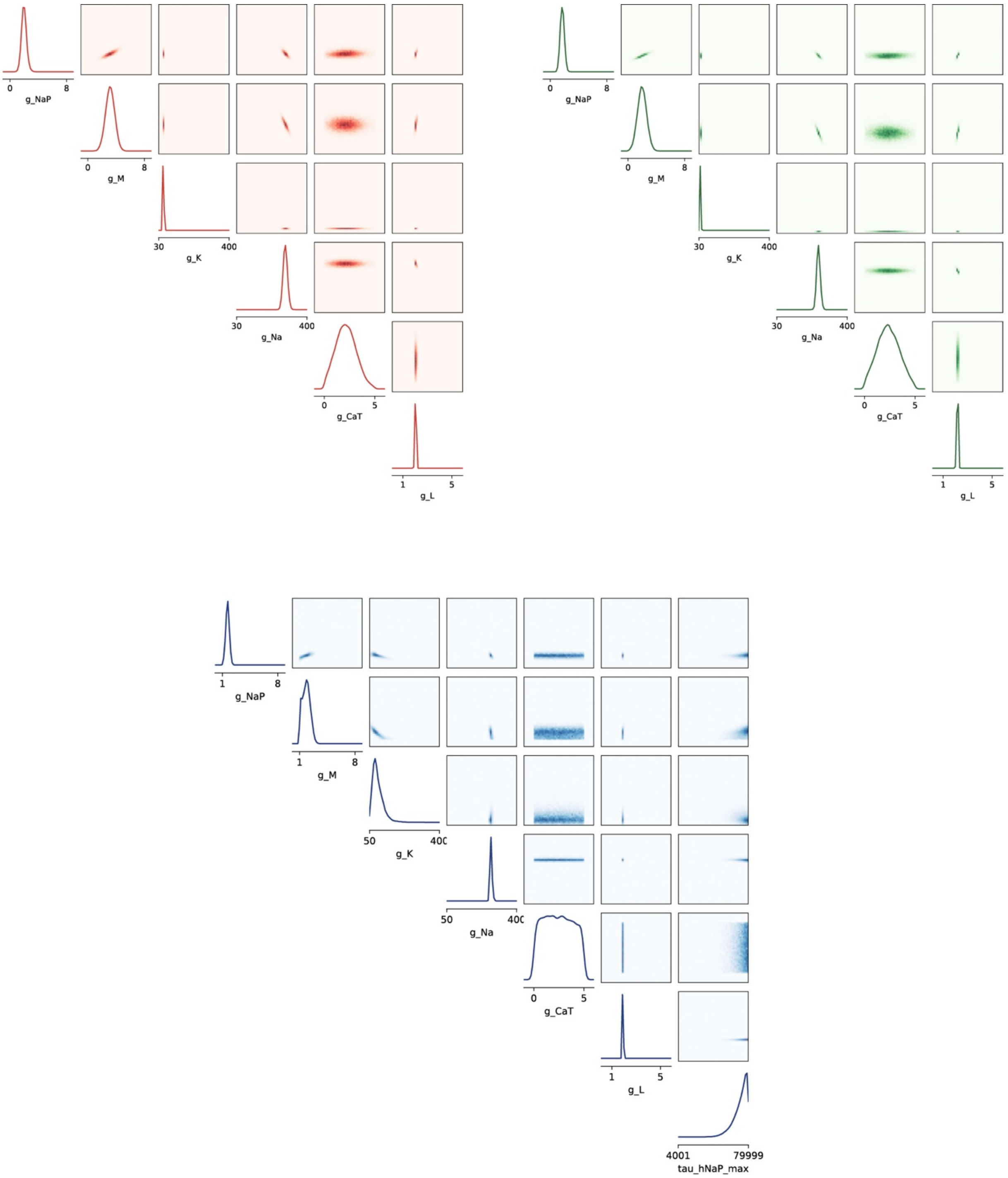


**Supplementary Figure 9. Pairwise posterior distributions of inferred conductances.** Paired plots showing the marginal (diagonal) and bivariate (off-diagonal) posterior distributions of the estimated conductances for Cluster 1 (red, top left), Cluster 2 (green, top right), and Cluster 3 (blue, bottom). Off-diagonal panels display the joint distribution of each parameter pair, revealing dependencies between conductances in reproducing each burst phenotype. For Cluster 3, an additional parameter controlling $I_{NaP}$ deinactivation kinetics ($\tau_{hNaPmax}$) is included as a seventh estimated parameter (rightmost column and bottom row).

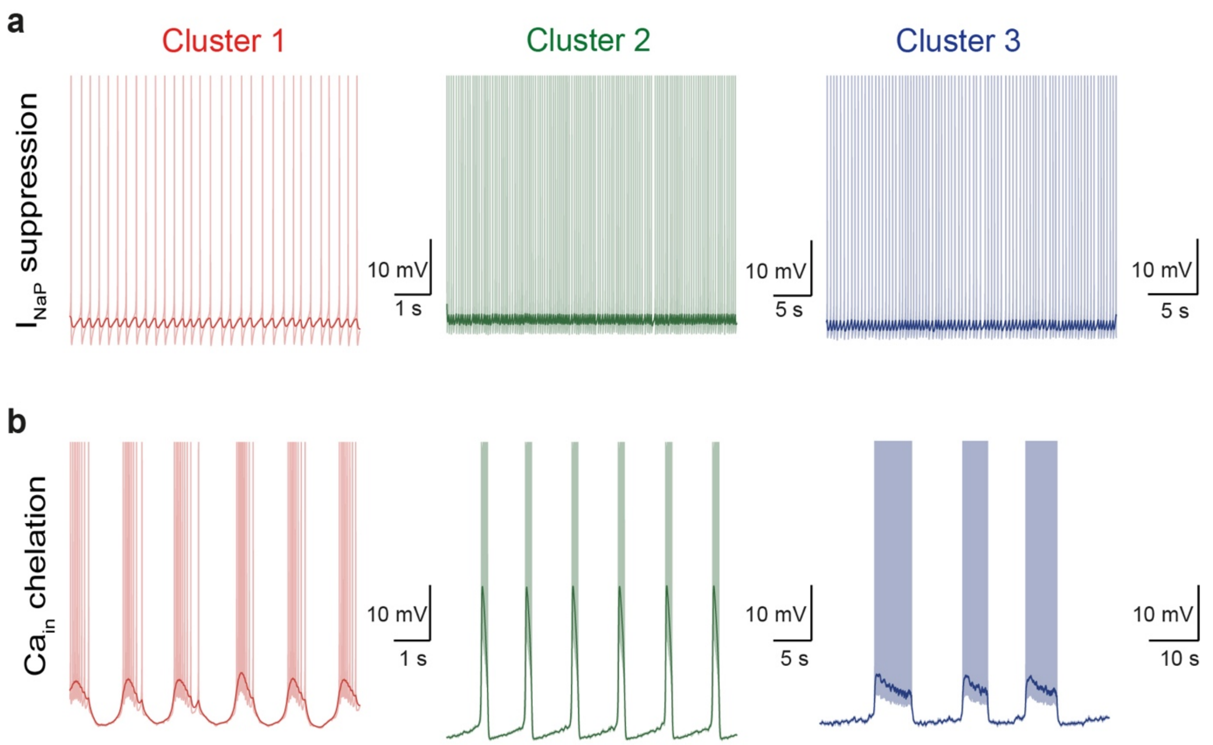


**Supplementary Figure 10. Additional *in silico* pharmacological manipulations applied to the fitted models.** **a** Effect of persistent sodium current suppression ($g_{NaP}$ = 0) under apamin-mimicking conditions ($g_{SK}$ = 0; cluster-specific $I_{inj}$ values in Table 2). **b** Effect of intracellular calcium chelation (kchelation = 0.05) in the presence of SK conductance (cluster-specific values in Table 2). Light-colored curves show individual simulation traces; thick dark-colored curves indicate the burst envelope
